# A fold switch regulates conformation of an alphavirus virus RNA-dependent RNA polymerase

**DOI:** 10.1101/2025.09.18.676892

**Authors:** Jamie J. Arnold, Sean M. Braet, Luiz C. Vieira, Ibrahim M. Moustafa, David W. Gohara, Julia A. Fecko, Yuan-Wei Norman Su, Abha Jain, David Aponte-Diaz, Claus O. Wilke, Ganesh S. Anand, Neela H. Yennawar, Craig E. Cameron

## Abstract

Alphaviruses are mosquito-vectored, positive-strand RNA viruses causing rheumatic and neurological diseases. Like all RNA viruses, they encode an RNA-dependent RNA polymerase (RdRp, nsP4). Purification of an nsP4 derivative capable of processive RNA synthesis from a heteropolymeric template has been unsuccessful. Prior studies indicated O’nyong-nyong virus (ONNV) nsP4 is soluble and requires additional non-structural proteins for activity. We performed biochemical and biophysical characterization of ONNV nsP4, including analytical ultracentrifugation and small-angle X-ray scattering (SAXS), revealing an extended conformation inconsistent with AlphaFold predictions of a compact structure. Fold switching was required for the extended conformation. Hydrogen-deuterium exchange mass spectrometry confirmed the fold-switched, extended state. Phylogenetic analysis showed conservation of residues contributing to both extended and compact states, implying functional roles for each. The extended form exhibited weak RNA binding and no polymerase activity on primed templates. The SAXS envelope of a precursor containing 50 amino acids from the nsP3 C-terminus (CT50-P34) matched the compact state. We propose precursor forms adopt the compact conformation. At the replication site, proteolytic cleavage would convert the precursor to an active polymerase. Polymerase dissociation upon completion of synthesis would induce fold switching to the inactive, extended state, precluding cytoplasmic activity that would activate intracellular immune responses.

## INTRODUCTION

The viral RNA-dependent RNA polymerase (RdRp) is a superb target for therapeutic control of infections by RNA viruses (1–3). Ribavirin (4), sofosbuvir (5,6), favipiravir (7,8), remdesivir (9,10), and molnupiravir (11) are all nucleoside analogs exhibiting broad-spectrum antiviral activity in a clinical setting. Most antiviral nucleosides are discovered by screening libraries of compounds for antiviral activity, often by using a biological assay for screening. For a nucleoside analog to exhibit antiviral activity in cells, the molecule must be converted to the active, triphosphorylated form. As a result, nucleoside analogs with the potential to interfere with virus multiplication will be missed when the analog cannot be converted to the active metabolite. It is now feasible to create nucleoside prodrugs capable of bypassing blocks to intracellular metabolism (12,13). The ability to use purified RdRp-based assays for screening offers an alternative pathway to discovery of efficacious antiviral nucleotide analogs. Access to purified RdRp is also essential to elucidating mechanism of action of antiviral nucleotides.

Alphaviruses are arthropod-borne, positive-strand RNA viruses belonging to the Togavirus family of viruses. The alphavirus genus has been divided into two major groups: old world and new world viruses (14,15). Old world viruses include Chikungunya virus (CHIKV) and O’nyong-nyong virus (ONNV), among many others (14–17). New world viruses include all of the equine encephalitis viruses (14,15,18). Infection by both groups of viruses causes substantial inflammation of the joints (old world) or the central nervous system (new world). Alphaviruses are prominent members of any list of RNA viruses with pandemic potential (19).

Western equine encephalitis virus was the first alphavirus discovered, and this discovery was made nearly 100 years ago (20). Since that time, the significance of these viruses in terms of human infectious disease has continued to increase and many more alphaviruses discovered. Surprisingly, not a single, active preparation of a purified alphavirus RdRp has been reported. To clarify, *active* here refers to the ability to initiate and/or to elongate on a heteropolymeric RNA template for multiple cycles of nucleotide incorporation. Terminal transferase activity of an alphavirus RdRp has been observed (21,22). Also, extension of a hairpin RNA containing an oligo(rU) has been reported (23). None of the substrates used in these previous studies permit a robust, rigorous, mechanistic analysis of the alphavirus RdRp.

The alphavirus nsP4-coding sequence encodes an amino-terminal domain of 100-160 amino acids, depending on the study considered, that is not found in other related RdRps (22–25). The response of many investigators to this substantive difference between RdRps has been to delete the unique domain (22,23,25). The hypothesis that *extra* residues, when compared to related enzymes, are not essential for function may not be sound. Elaboration of a known enzymatic domain with additional domains could confer better regulation of the enzymatic activity relative to enzymes lacking the domain. In this scenario, for such a regulatory domain to be essential for overall function would not be surprising.

Inspired by studies of others using Sindbis and Ross River viruses (23), we have performed a comprehensive biochemical and biophysical analysis of full-length ONNV RdRp. The protein was monomeric and well behaved in solution. A combination of circular dichroism spectroscopy, size exclusion chromatography, multi-angle light scattering, analytical ultracentrifugation, small-angle X-ray scattering, and amide hydrogen-deuterium exchange mass spectrometry were all consistent with the full-length enzyme adopting an elongated conformation that does not reflect the typical, globular, “right-hand” conformation of a viral RdRp (26) or the conformation of nsP4 predicted by AlphaFold (24). The extended conformation was unable to bind to RNA with high affinity and unable to support primed extension on any substrate tested.

Alphaviruses encode four non-structural proteins: nsP1, nsP2, nsP3, and nsP4 produced from two different precursors: P123 and P1234 (14,27–29). Processing of P1234 in Sindbis virus yields two smaller precursors P12 and P34 (30). Lemm and Rice showed that the combination of P123 and P34 were sufficient for genome replication (31). Curiously, a P34 derivative containing only carboxy-terminal 50 amino acids of nsP3 linked to nsP4 (CT50-P34) was able to fully substitute for P34 (32). We have purified CT50-P34. The SAXS envelope of this protein was consistent with a tetramer of CT50-P34 in the compact conformation.

Together, these observations suggest that nsP4 can exist in two distinct conformations. We hypothesize that a fold switch of the compact, presumably active conformation, produces the extended, inactive conformation reported here. We suggest that the fold switch from compact to extended prevents aberrant activity of ONNV RdRp when not engaged in genome replication, perhaps to prevent formation of double-stranded RNA in the cytoplasm. Importantly, key interfaces stabilizing the extended conformation are conserved across the entire alphavirus genus consistent with all alphavirus polymerases exhibiting similar regulation. Finally, we discuss the potential utility of CT50-P34 as a model for fold-switching proteins broadly.

## MATERIALS AND METHODS

### Materials

DNA oligonucleotides and dsDNA fragments, GBlocks, were from Integrated DNA Technologies. RNA oligonucleotides were either from Horizon Discovery Ltd. (Dharmacon) or Integrated DNA Technologies. Restriction enzymes were from New England Biolabs. IN-FUSION HD enzyme was from TakaraBio. T4 polynucleotide kinase was from ThermoFisher. Nuvia Ni-NTA, UNOsphere Q, UNOsphere S pre-packed affinity columns were from Bio-Rad. [γ-^32^P]ATP (6,000 Ci/mmol) and [α-^32^P]ATP (3,000 Ci/mmol) were from Perkin Elmer. Nucleoside 5’-triphosphates (ultrapure solutions) were from Cytiva. All other reagents were of the highest grade available from MilliporeSigma, VWR, or Fisher Scientific.

### Construction of modified pSUMO vectors containing TWIN-STREP-Tag

The pSUMO system allows for production of SUMO fusion proteins containing an amino-terminal affinity tag fused to SUMO that can be purified by affinity chromatography and subsequently processed by the SUMO protease, Ulp1 (33). After cleavage this will produce an authentic untagged protein target of interest. The pSUMO vector (LifeSensors) (33)was modified such that the amino terminal coding sequence encoded for a TWINS-TREP-tag and a six-histidine tag, both codon optimized for bacterial expression. The DNA sequences (GBlocks encoding the tags) were cloned into pSUMO using XbaI and SalI by IN-FUSION. The final construct (pTWIN-STREP-6HIS-SUMO) were confirmed by sanger sequencing performed by Genewiz.

### Codon-optimized sequence for TWIN-STREP-6HIS-tag

5’atgggaagttggtcacacccccaatttgaaaagggcggttccggctcgagctggagccatccgcagttcgagaaaggttctagcggtcatcatcaccaccaccacggctccagcggt3’ (amino acid sequence: MGSWSHPQFEKGGSGSSWSHPQFEKGSSGHHHHHHGSSG)

### Construction of ONNV nsP4 and CT50-P34 bacterial expression plasmids

The ONNV nsP4 gene was codon optimized for expression in *E. coli* and obtained from Integrated DNA Technologies. The amino acid sequence for nsP4 was derived from ONNV (GenBank AAA46784.1). The gene (IDT GBlock) was directly cloned into the pTWIN-STREP-6HIS-pSUMO bacterial expression plasmid using BsaI and SalI. The amino acid sequence for the last 50 amino acids of nsP3 was codon optimized for expression in E coli. The Opal stop codon was replaced with an arginine codon. The CT50-P34 was cloned into the pET26Ub-CHIS bacterial expression plasmid using SacII and BamHI sites (34). The final constructs were confirmed by sanger sequencing performed by Genewiz.

Codon optimized sequence for ONNV nsP4 –5’tatatatttagttcagatacaggacaagggcacctgcagcaaaaaagcgttcgtcagacaaccttgccggttaacattgttgaggaagtgcacgaagagaagtgctatccaccgaagctggacgagattaaagaacaactgttgctcaagcgcctgcaagagagcgcgtctaccgcgaaccgtagccgttatcaaagcagaaaagtcgaaaacatgaaagcgaccatcatccaccgccttaaagaaggttgtcgtctttacctcgcgtctgaaacgccgcgtgtgccctcttaccgcgtgacttatccggcaccgatttattccccgtccattaacatcaaactgactaatccggaaactgctgtcgcagtctgtaatgaattcctggctcgcaactaccctactgtagcgagctaccaggtgacggacgagtacgacgcttacctggatatggttgacggcagcgagagctgcctggaccgcgcgaccttcaacccgtcgaagctcagatcttacccgaaacagcatagctaccatgcaccaaccatccgttctgcggtgccgtccccgtttcagaacacgctgcaaaacgtcttagcggcggctaccaagcgtaattgtaacgtgacgcagatgcgtgaactgccaaccatggacagcgccgtttttaacgtcgagtgctttaagaagtacgcgtgcaatcaggagtactggcgtgagttcgcatcgagcccgattcgtgtcacgaccgagaatctcaccatgtatgttacgaaattaaagggtccgaaggccgccgctctgttcgcgaaaacccataatctgttgccgctgcaagaggtgccgatggatcgtttcaccatggacatgaaacgcgacgttaaagttaccccgggtactaagcacactgaggaacgtccgaaggtgcaggtgatccaggcagcggagccgctggcgaccgcatatctgtgtggcattcaccgtgagttagttcgtcgcttgaatgcggtgctgctgccaaatgtgcacaccttattcgacatgtccgcggaagattttgacgcgattatcgccacccactttaagccgggtgacgccgtactggaaaccgatattgcttcatttgataaatcgcaagatgatagcctggcgtccaccgctatgatgctgctggaggacctgggtgtggaccaaccgatcctggacctgattgaagcagcgttcggcgaaatcagcagctgccatctgccgaccggtacgcgtttcaagttcggcgcaatgatgaagagtggtatgtttttgaccctgttcgtgaataccctgctgaacataacgatcgcgtctagggtcctggaagagagattgaccacctccgcctgcgcagcgttcatcggcgacgataacattattcacggcgttgttagcgatgccctgatggccgcgcgttgcgccacctggatgaacatggaagtgaaaatcatcgatgcggtggttagcgaaaaagcgccttatttctgcggtggttttatcctccatgataccgtcaccggcacttcctgccgtgtcgctgatccgttgaagcgcctgttcaagcttggcaaaccgctggctgctggtgacgagcaagacgaggatcgccgtcgtgccctcgcggatgaggtgacccgttggcagcgtaccggtctggttacggagttggaaaaagcggtttatagccgttatgaagtacagggcatcaccgcggttattaccagcatggcaacctttgcgaacagcaaagagaactttaaaaaattgcgcggtccggtggttaccttgtacggcggtccgaag3’

### Expression of ONNV nsP4

*E. coli* BL21(DE3) competent cells were transformed with the pTWIN-STREP-6HIS-SUMO-ONNV-nsP4 plasmid for protein expression. BL21(DE3) cells containing the pTWIN-STREP-6HIS-SUMO-ONNV-nsP4 plasmid were grown in 100 mL of media (NZCYM) supplemented with kanamycin (K25, 25 µg/mL) at 37 °C until an OD_600_ of 1.0 was reached. This culture was then used to inoculate 4L of K75-supplemented ZYP-5052 auto-induction media at 37 °C. The cells were grown at 37 °C to an OD_600_ of 0.8 to 1.0, cooled to 15 °C and then grown for 36-44 h. After ∼40 h at 15 °C the OD_600_ reached ∼10–15. Induction was verified by SDS-PAGE using TGX-stain free gels and visualized by using Bio-Rad ChemiDoc MP Imager. Cells were harvested by centrifugation (6000 x *g*, 10 min) and the cell pellet was washed once in 200 mL of TE buffer (10 mM Tris, 1 mM EDTA), centrifuged again, and the cell paste weighed.

### Expression of ONNV CT50-P34-CHIS

*E. coli* BL21(DE3)pCG1 competent cells were transformed with the pET26Ub-ONNV-CT50_P34-CHIS plasmid for protein expression. BL21(DE3)pCG1 cells containing the pET26Ub-ONNV-CT50-P34-CHIS plasmid were grown in 100 mL of media (NZCYM) supplemented with kanamycin (K25, 25 µg/mL) and chloramphenicol (C20, 20 µg/mL) at 37 °C until an OD_600_ of 1.0 was reached. This culture was then used to inoculate 4L of C60,K75-supplemented ZYP-5052 auto-induction media at 37 °C. The cells were grown at 37 °C to an OD_600_ of 0.8 to 1.0, cooled to 15 °C and then grown for 36-44 h. After ∼40 h at 15 °C the OD_600_ reached ∼10–15. Induction was verified by SDS-PAGE using TGX-stain free gels and visualized by using Bio-Rad ChemiDoc MP Imager. Cells were harvested by centrifugation (6000 x *g*, 10 min) and the cell pellet was washed once in 200 mL of TE buffer (10 mM Tris, 1 mM EDTA), centrifuged again, and the cell paste weighed.

### Purification of ONNV nsP4

Frozen cell pellets were thawed on ice and suspended in lysis buffer (100 mM Potassium Phosphate pH 8.0, 500 mM NaCl, 2 mM TCEP, 20% glycerol, 1.4 μg/mL leupeptin, 1.0 μg/mL pepstatin A and two Roche EDTA-free protease tablet per 5 g cell pellet), with 5 mL of lysis buffer per 1 gram of cells. The cell suspension was lysed by passing through a Nano DeBEE Gen II Homogenizer using nozzle Z08 in reverse flow mode (BEE International) at 20,000 psi. After lysis, phenylmethylsulfonylfluoride (PMSF) and NP-40 were added to a final concentration of 1 mM and 0.1% (v/v), respectively. While stirring the lysate, polyethylenimine (PEI) was slowly added to a final concentration of 0.25% (v/v) to precipitate nucleic acids from cell extracts. The lysate was stirred for an additional 30 min at 4 °C after the last addition of PEI, and then centrifuged at 75,000 x g for 30 min at 4 °C. The PEI supernatant was decanted and solid ammonium sulfate was slowly added to 40% saturation while stirring. The granular ammonium sulfate was pulverized to a fine powder before use by using a mortar and pestle. The solution was stirred for an additional 15 - 30 min at 4 °C after addition of ammonium sulfate. The ammonium sulfate precipitated material was pelleted by centrifugation for 30 min at 25,000 rpm (75,000 x *g*) at 4 °C. The supernatant was decanted and the pellet was suspended in lysis buffer. The resuspended ammonium sulfate pellet was then loaded onto a 5 mL Nuvia IMAC Ni column (Bio-Rad) at a flow rate of 1 mL/min (approximately 1 mL bed volume per 100 mg total protein) equilibrated with buffer A (25 mM HEPES pH 7.5, 500 mM NaCl, 2 mM TCEP, 20% glycerol) with 5 mM imidazole using a Bio-Rad NGC System. After loading, the column was washed with twenty column volumes of buffer A with 5 mM imidazole and 5 column volumes of buffer A with 50 mM imidazole. The protein was eluted using buffer A with 500 mM imidazole. Ulp1 (1 µg per mg) was added to the eluted protein to cleave the SUMO fusion and was dialyzed overnight against 1 L buffer B (50 mM HEPES pH 7.5, 2 mM TCEP, 20% glycerol) containing 500 mM NaCl (membrane employed a 12 −14,000 Da MWCO).

After dialysis the dialyzed sample was diluted in Buffer A to a final salt concentration of 50 mM. The sample was loaded onto a 1 mL UNOsphere Q column in tandem with a 1 mL UNOsphere S column (1 mL bed volume/25 mg of protein) at 1 mL/min. The protein passes through the Q column and binds to the S column. Both columns were washed with two column volumes of Buffer B containing 50 mM NaCl, the S column was subsequently washed with 10 column volumes of Buffer B containing 50 mM NaCl and the protein eluted form the S column using a linear gradient (6 column volumes) from 50 to 1000 mM NaCl in Buffer B. The eluted protein was pooled and then passed through a Cytiva HiLoad Superdex 16/600 200 pg size exclusion column equilibrated with Buffer B containing 500 mM NaCl. nsP4 fractions were pooled, concentrated, and dialyzed against 25 mM HEPES pH 7.5, 1 mM TCEP, 5% glycerol, 500 mM NaCl. The protein concentration was determined by measuring the absorbance at 280 nm by using a Nanodrop spectrophotometer and using a calculated molar extinction coefficient of 47,790 M^-1^ cm^-1^. Purified, concentrated protein was aliquoted and frozen at −80 °C until use. Typical nsP4 yields were 1 mg/10 g of *E. coli* cells.

### Purification of ONNV CT50-P34-CHIS

Frozen cell pellets were thawed on ice and suspended in lysis buffer (100 mM Potassium Phosphate pH 8.0, 500 mM NaCl, 2 mM TCEP, 20% glycerol, 1.4 μg/mL leupeptin, 1.0 μg/mL pepstatin A and two Roche EDTA-free protease tablet per 5 g cell pellet), with 5 mL of lysis buffer per 1 gram of cells. The cell suspension was lysed by passing through a Nano DeBEE Gen II Homogenizer using nozzle Z08 in reverse flow mode (BEE International) at 20,000 psi. After lysis, phenylmethylsulfonylfluoride (PMSF) and NP-40 were added to a final concentration of 1 mM and 0.1% (v/v), respectively. While stirring the lysate, polyethylenimine (PEI) was slowly added to a final concentration of 0.25% (v/v) to precipitate nucleic acids from cell extracts. The lysate was stirred for an additional 30 min at 4 °C after the last addition of PEI, and then centrifuged at 75,000 x g for 30 min at 4 °C. The PEI supernatant was decanted and solid ammonium sulfate was slowly added to 40% saturation while stirring. The granular ammonium sulfate was pulverized to a fine powder before use by using a mortar and pestle. The solution was stirred for an additional 15 - 30 min at 4 °C after addition of ammonium sulfate. The ammonium sulfate precipitated material was pelleted by centrifugation for 30 min at 25,000 rpm (75,000 x *g*) at 4 °C. The supernatant was decanted and the pellet was suspended in lysis buffer. The resuspended ammonium sulfate pellet was then loaded onto a 5 mL Nuvia IMAC Ni column (Bio-Rad) at a flow rate of 1 mL/min (approximately 1 mL bed volume per 100 mg total protein) equilibrated with buffer A (25 mM HEPES pH 7.5, 500 mM NaCl, 2 mM TCEP, 20% glycerol) with 5 mM imidazole using a Bio-Rad NGC System. After loading, the column was washed with twenty column volumes of buffer A with 5 mM imidazole and 5 column volumes of buffer A with 50 mM imidazole.

The protein was eluted using buffer A with 500 mM imidazole. The eluted protein was dialyzed overnight against 1 L buffer B (50 mM HEPES pH 7.5, 2 mM TCEP, 20% glycerol) containing 500 mM NaCl (membrane employed a 12 −14,000 Da MWCO). After dialysis the dialyzed sample then passed through a Cytiva HiLoad Superdex 16/600 200 pg size exclusion column equilibrated with Buffer B containing 500 mM NaCl. CT50_P34-CHIS fractions were pooled and concentrated. Protein was passed through two consecutive Zeba desalting columns equilibrated with 25 mM HEPES pH 7.5, 1 mM TCEP, 5% glycerol, 500 mM NaCl. The protein concentration was determined by measuring the absorbance at 280 nm by using a Nanodrop spectrophotometer and using a calculated molar extinction coefficient of 53,290 M^-1^ cm^-1^. Purified, concentrated protein was aliquoted and frozen at −80 °C until use. Typical CT50-P34-CHIS yields were 1 mg/10 g of *E. coli* cells.

### Expression and purification of Enterovirus and Flavivirus RdRp

Poliovirus 3Dpol RdRp and Zika virus NS5 RdRp were expressed and purified essentially as described previously (33–35).

### 5’-32P-labeling of RNA substrates

RNA oligonucleotides were end-labeled by using [*γ*- ^32^P]ATP and T4 polynucleotide kinase. Reaction mixtures, with a typical volume of 50 *μ*L, contained 0.5 *μ*M [*γ*-^32^P]ATP, 10 *μ*M RNA oligonucleotide, 1× kinase buffer, and 0.4 unit/*μ*L T4 polynucleotide kinase. Reaction mixtures were incubated at 37 °C for 60 min and then held at 65 °C for 5 min to heat inactivate T4 PNK. For RNAs containing a 3’phosphate T4 PNK (minus 3’phosphatase) was used using the same reaction conditions.

### Annealing of dsRNA substrates

dsRNA substrates were produced by annealing 10 μM RNA oligonucleotides in T_10_E_1_ [10 mM Tris pH 8.0 and 1 mM EDTA] and 50 mM NaCl in a Progene Thermocycler (Techne). Annealing reaction mixtures were heated to 90 °C for 1 min and either snap cooled on ice or slowly cooled (5 °C/min) to 10 °C. Specific scaffolds are described in the figure legends.

### RNA-dependent RNA polymerase assays

Reactions contained 1 µM 32-P-labeled RNA primer/template (T1H or sym/sub-U), 2 µM RdRp, 1 mM ATP or 1 mM NTPs. Additionally, reactions contained 10 µM pGGC trinucleotide primer, 1 µM RNA template, 10 µM NTPs and 0.1 µCi/µL alpha32P-ATP. Reactions were performed in 25 mM HEPES pH 7.5, 5 mM MgAcetate, 1 mM TCEP and 20 mM Potassium Glutamate. Reactions were initiated by addition of RdRp and incubated at 30 oC for 30 min. Reactions were quenched by addition of 50 mM EDTA.

### Denaturing PAGE analysis

An equal volume of loading buffer (85% formamide, 0.025% bromophenol blue and 0.025% xylene cyanol) was added to quenched reaction mixtures and heated to 90 °C for 5 min prior to loading 5 µL on a denaturing either 15% or 23% polyacrylamide gel containing 1X TBE (89 mM Tris base, 89 mM boric acid, and 2 mM EDTA) and 7 M urea. For reactions that contained dsRNA substrates an excess (50-fold) of unlabeled RNA oligonucleotide (trap strand) that is the exact same sequence to the ^32^P-labeled RNA oligonucleotide in the reaction was present in the loading buffer to ensure complete separation and release of the ^32^P-lableled RNA oligonucleotide prior to gel electrophoresis (36). This procedure allows efficient strand separation of ^32^P-labeled RNA oligos that are in the presence of their RNA complements (36). Electrophoresis was performed in 1x TBE at 90 W. Gels were visualized by using a PhosphorImager (GE) and quantified by using ImageQuant TL software (GE).

### RNA binding assays

To test interactions between RdRp’s and RNA, increasing concentrations of RdRp were added into a solution containing 10 nM of fluorescein-labeled nucleic acid in a binding buffer [25 mM HEPES pH 7.5, 5 mM magnesium acetate, 1 mM TCEP, and 20 mM Potassium Glutamate] in a 30 μL final reaction volume. Reactions were incubated for 5 min at 30 °C and then measured. Experiments were performed in a 384-well plate format using a BioTek H1M1 plate reader using a dual wavelength 485/20 485/20 fluorescence polarization cube. Data from protein titration experiments were fit to a hyperbola:

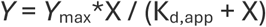

where *X* is the concentration of protein, *Y* is degree of polarization, *K*_d,app_ is the apparent dissociation constant, and *Y*_max_ is the maximum value of *Y*.

### Circular Dichroism Spectroscopy

Circular dichroism (CD) spectroscopy measurements were conducted using a JASCO J-1500 spectrometer, featuring a Peltier model PTC-517 thermostat cell holder. Signals were recorded within the range of 260 nm to 185 nm, employing a scan speed of 50 nm/min and a band width of 1 nm, all maintained at 20°C. Throughout the experiments, the data pitch was set to 1 nm, and the digital integration time (DIT) was 4 seconds. The pathlength of the quartz cell used was 1 mm, and the concentration of nsP4 was 0.29 mg/mL. Prior to sample measurement, a buffer blank was run for baseline subtraction, and the resulting data were converted to units of molar ellipticity. Data analysis was conducted using both the Jasco Multivariate SSE program and BestSel single spectra analysis and fold recognition software. Additionally, the predicted CD values were obtained using PDB2CD software.

### Size Exclusion Chromatography Multi-Angle Light Scattering (SEC-MALS)

The SEC-MALS experiment utilized an Agilent 1260 Infinity II HPLC system with an autosampler. A Wyatt SEC hydrophilic column featuring 5-µm silica beads, 100 Å pore size, and dimensions of 7.8 × 300 mm was employed. Analysis of the molar mass of eluted peaks was conducted using Wyatt Technology’s DAWN MALS and Optilab refractive index detector, alongside an Agilent UV detector. Prior to sample injection, the SEC-MALS system was equilibrated for 5 hours with a buffer comprising 25mM Hepes (pH 7.5), 500mM NaCl, 1mM TCEP, and 5% glycerol. UV detection was set at 280nm, and the temperature was maintained at 4°C. Injection of 50µl of Nsp4 at 2.9mg/ml was followed by column operation at a flow rate of 0.5 ml/min, with a total chromatogram run time of 40 minutes. Normalization and alignment of the MALS and refractive index detectors were performed using a standard BSA run in the same buffer prior to the sample injection. UV served as the concentration source. Data analysis was conducted using ASTRA software, version 8.0.2 (Wyatt). The chromatogram exhibited a single, monodisperse peak in both light scattering and refractive index detection. The molar mass determination indicated a monomer size of 69.7 kDa.

### Analytical Ultracentrifugation (AUC)

The AUC analysis was conducted using an Optima multiwavelength instrument from Beckman Coulter, Indianapolis, IN, equipped with absorbance and interference optics. The instrument operated under full vacuum conditions with the rotor spinning at 40,000 RPM for 16.5 hours at 5°C. Radial scans of the cells were captured every 63 seconds at 280 nm. AUC data was processed using UltraScan III. Reference scans were manually chosen to convert raw radial intensity data to pseudo-absorbance. The air-liquid meniscus was manually determined for each sector, and the data range was manually set between approximately 6.0 cm and 7.1 cm. Initial scans, typically the first five to 10, lacking significant sedimentation signal, were excluded from analysis. Similarly, scans at the end with minimal sedimentation signal were omitted. The data was fitted with an S-value range of 1 to 10 and a resolution of 100. The frictional ratio range was set between 1 and 4 with a resolution of 64. Time-invariant noise was addressed during the initial two-dimensional spectral analysis. When residuals fell below 0.003, the data underwent re-fitting for time and radially invariant noise, and adjustments were made to the meniscus position. Upon identifying the correct meniscus, an additional fitting for time and radial invariant noise was executed using an iterative method with a maximum of 10 iterations. Finally, 64 Monte Carlo simulations were conducted, and the resulting pseudo-three-dimensional plots were analyzed to determine the final S-values, frictional ratios, and molecular weight calculations.

### Small-Angle X-ray Scattering (SAXS)

BioSAXS experiments were conducted on nsP4 protein, at concentrations of 2.9 mg/ml and 1.0 mg/ml. The experiments were performed in 25 mM HEPES pH 7.5, 1 mM TCEP, 5% glycerol and 500 mM NaCl. X-rays for BioSAXS data acquisition were generated by a Rigaku MM007 rotating anode X-ray source, coupled with the BioSAXS2000nano Kratky camera system. This system incorporates OptiSAXS confocal max-flux optics and a HyPix-3000 Hybrid Photon Counting detector. The sample-to-detector distance was set at 495.5 mm, calibrated using silver behenate powder. The usable momentum transfer scattering vector (q-space) ranged from qmin= 0.008 Å^-1^ to qmax= 0.6 Å^-1^. The X-ray beam energy was 1.2 keV, with a Kratky block attenuation of 22% and a beam diameter of approximately 100 μm. Protein samples were introduced using the Rigaku autosampler into a quartz capillary flow cell cooled to 4°C. The entire X-ray flight path, including the beam stop, was maintained under vacuum conditions of < 1×10^-3^ torr to minimize air scatter. Automated data collection, featuring cleaning cycles between samples, was done using the Rigaku SAXSLAB software. Data processing within Rigaku SAXSLAB software included averaging of six ten-minute images and three replicates from both protein and buffer samples to ensure absence of X-ray radiation damage. SAXS data overlays confirmed no radiation decay or sample loss during the 60-minute data collection period and the replicate SAXS data overlaid well. Subsequently, reference buffer subtraction was applied to derive the raw SAXS scattering curve from protein alone. The forward scattering I(0) and the radius of gyration, Rg were computed using the Guinier approximation. Molecular mass estimation was facilitated by comparing the data to standard protein SAXS data for BSA. Analysis of data files included parameters such as Rg, maximum particle dimension, Dmax, Guinier fits, Kratky plots, and pair distance distribution function, P(r), using ATSAS software. High-q Kratky plots indicated well-folded proteins with no flexible regions. P(r) plots were calculated using GNOM, yielding the Dmax values. Solvent envelopes were generated using DAMMIF and DENSS algorithms. The best resolved structure of the Chikungunya virus replication complex structure where nsP4 is seen as a complex with the core RNA replicase nsP1 and the helicase-protease nsP2, was used for the initial SAXS fits (PDB code 7Y38). Manual fitting in Pymol of the fingers-domain of the NSP4 polymerase in the Denss electron density solvent envelope as derived from the SAXS data fits reasonably well. There is a conformational change seen in a 100-residue stretch of the N-terminal domain with respect to the fingers-domain. However the frictional ratio as from AUC data, Kratky plots from SAXS, fits between experimental and calculated SAXS profiles in the Crysol software as well circular dichroism data all point to a limited range of disorder for this segment with no significant changes in the secondary structures or unfolding of the two helices. Modeling of the N-terminal helices and analysis of the conformational change were done manually in Pymol and subsequently refined via normal mode analysis in Sreflex, utilizing the SAXS data. This refinement and a fit with a molecular dynamics simulated model both confirmed the 180° flip of the N-terminal helix leading to an extended inactive conformation for nsp4. Theoretical scattering profiles of the constructed model was computed and fitted to experimental scattering data using CRYSOL, resulting in good Chi-square fits.

### Amide Hydrogen Deuterium Exchange

Deuterium labeling buffer was prepared by diluting 10 X HEPES/Sodium Chloride (pH 7.5) in H_2_O in D_2_O (99.9%) (Final concentration: 500 mM NaCl, 25 mM HEPES, 1 mM TCEP). 3 μL of 1.1 mg/mL alphavirus nsP4 were added to 57 μL of labeling buffer for a final labeling concentration of 85.41%. Deuterium labeling was carried out for 1, 10, and 100 min at 20°C using a PAL-RTC (Leap) autosampler. After labeling, equivalent volumes of labelled sample and prechilled quench solution (1.5 M GdnHCl, 0.25 M TCEP) were added to bring the reaction to pH 2.5. Unlabeled reference data was also collected in triplicate by diluting 3 uL of 1.1 mg/mL alphavirus nsP4 in 500 mM NaCl, 25 mM HEPES, 1 mM TCEP in H_2_O (pH 7.5).

### Maximal Deuteration

Maximally deuterated nsP4 was prepared as described by Wales et al. (37). Briefly, 6 µL aliquots of aqueous nsP4 at 1 mg/mL were dessicated dried on a speed vacuum and resuspended in 3.5 uL 7 M GdnHCl and 50 mM DTT. 3 µL of resuspended sample was transferred to a PCR tube and heated to 90 °C for 5 min using a thermocycler. The sample was allowed to cool to 20 °C for 2 min prior to addition of 57 µL of labelling buffer. The labelling reaction was heated to 50 °C for 10 min using a thermocycler followed by cooling for 2 min at 20 °C and 0 °C. Chilled and labelled sample was mixed with an equal volume of prechilled quench solution prior to mass spectrometry analysis. Maximal deuteration was carried out across 4 technical replicates. Peptides in the maximally deuterated controls were only detected for the N-terminal 120 residues.

### Pepsin proteolysis and mass spectrometry analysis

Approximately 8–10 pmol of the sample were loaded onto a BEH pepsin column (2.1 × 30 mm) (Waters, Milford, MA) in 0.1% formic acid at 100 μL/min. Proteolyzed peptides were trapped in a 2.1 x 5 mm C18 trap (ACQUITY BEH C18 VanGuard Pre-column, 1.7 µM, Waters, Milford, MA). Peptides were eluted in an acetonitrile gradient (35%) in 0.1% formic acid on a reverse phase C18 column (AQUITY UPLC BEH C18 Column, Waters, Milford, MA) at 40 μL/min. All fluidics were controlled by nanoACQUITY Binary Solvent Manager (Waters, Milford, MA).

Electrospray ionization mode was utilized, and ionized peptides were sprayed onto an SYNAPT XS mass spectrometer (Waters, Milford, MA) acquired in HDMS^E^ Mode. Ion mobility settings of 600 m/s wave velocity and 197 m/s transfer wave velocity were used with collision energies of 4 and 2 V used for trap and transfer, respectively. High collision energy was ramped from 20 to 45 V while a 25 V cone voltage was used to obtain mass spectra ranging from 50 to 2000 Da (10 min) in positive ion mode. A flow rate of 5 µL/min was used to simultaneously inject 100 fmol/μL of [Glu^1^]-fibrinopeptide B ([Glu^1^]-Fib) as lockspray reference mass.

ProteinLynx Global Server (PLGS v 3.0, Waters, Milford, MA) was used to identify nsP4 peptides from reference undeuterated spectra. Deuterium uptake was assessed using DynamX v3.0 (Waters, Milford, MA) by comparing centroids of labelled mass spectra to unlabeled reference spectra. Peptides were subject to additional filters of: minimum intensity = 2000, minimum peptide length = 4, maximum peptide length = 25, minimum products per amino acid = 0.2, and precursor ion error tolerance <10 ppm, followed by manual curation based on spectral quality. Relative deuterium exchange plots and heatmaps were generated by DynamX v3.0. Cluster files were exported from DynamX v3.0 and analyzed using Deuteros 2.0 (38). Back exchange correction was applied using uptake values for maximally deuterated peptides or average back exchange from maximum deuteration replicates for peptides that were not detected in maximally deuterated samples. HDXMS datasets will be deposited to the ProteomeXchange Consortium via the PRIDE partner repository.

### Sequence Conservation Analysis of Alphavirus nsP4 Protein

To assess the conservation of the nsP4 protein across various alphaviruses, we conducted an effective number analysis. The process began with the acquisition of reference sequences for alphavirus nsP4 proteins (39). These reference sequences were then subjected to a Psi-BLAST search against the Protein Reference Viral DataBase (RVDB) (40) to retrieve a more diverse set of nsP4 protein sequences, enhancing the sequence diversity of the analysis.

Following the sequence retrieval, we calculated the entropy for each position in the multiple sequence alignment (MSA) to measure the variability across the aligned sequences. Entropy serves as an indicator of sequence conservation, with lower entropy values reflecting higher conservation. To further quantify sequence conservation, we computed the effective number (Neff) of sequences at each position in the alignment. Neff provides a measure of the diversity of the sequences contributing to each position, with higher Neff values indicating lower conservation.

Finally, using a custom Python script (41), we visualized the sequence alignment and corresponding conservation data. The alignment plot was generated to display the conservation levels across the nsP4 protein, with positions color-coded according to their Neff values. This plot allows for the identification of highly conserved regions within the nsP4 protein, which may be critical for its function.

### Direct Coupling Analysis (DCA)

To investigate the coevolutionary relationships between residues in the nsP4 protein of alphaviruses, we performed Direct Coupling Analysis (DCA) using the pyDCA library (42). The analysis began with the nsP4 multiple sequence alignment (MSA) described in section “<u>Sequence Conservation Analysis of Alphavirus nsP4</u> <u>Protein</u>”. The MSA was then processed using pyDCA’s mean-field DCA (mfDCA) algorithm to calculate the pairwise couplings between residues, which reflect the coevolutionary signals. The DCA results were expressed as coupling scores, with higher scores indicating stronger coevolutionary relationships. To enhance the interpretation, we applied the Average Product Correction (APC) to the raw coupling scores and further converted them into Z-scores. Residue pairs with Z-scores exceeding a defined threshold (>2) (corresponding to the ∼95th percentile) were selected as candidates with potentially meaningful covariation that may indicate functional or structural interactions. These significant pairs were then analyzed in the context of the protein’s structure to identify regions of interest.

### Relative Solvent Accessibility (RSA) and Residue Distance Calculations

To analyze the structural features of the nsP4 protein, we calculated the relative solvent accessibility (RSA) and the distances between specific residue pairs using the Biopython library (43).

*RSA Calculation:* The RSA values were obtained using the DSSP module of Biopython, which computes the solvent accessibility of each residue within a protein structure. We first parsed the protein structure file using the PDBParser from Biopython and then applied the DSSP algorithm to the nsP4 protein model. DSSP calculates the absolute solvent accessibility (ASA) for each residue and normalizes it against the maximum possible ASA for that residue type, resulting in the RSA value. Residues with higher RSA values are more exposed to the solvent, indicating that they are located on the surface of the protein.

*Distance Calculation:* To explore the spatial relationships between residues, we calculated the distances between the Cα atoms of residue pairs within the nsP4 protein. The distances were computed using Biopython’s NeighborSearch module. We selected all residues with Cα atoms and calculated the Euclidean distance between each pair of residues.

### Visualization of Solvent Accessibility and Residue Pair Interactions in nsP4 Protein

To visually represent the structural features of the nsP4 protein, we utilized PyMOL (44), a molecular visualization tool, to create detailed structural figures. Our approach involved modifying the B-factor field in the PDB file to encode relative solvent accessibility (RSA) values, which allowed us to color-code the structure based on solvent exposure.

*Modification of B-factor Values:* After calculating the RSA values for each residue using the DSSP module in Biopython, we replaced the B-factor values in the original PDB structure with these RSA values. This modification was necessary because PyMOL allows the coloring of structures based on B-factor values, providing a straightforward method to visualize solvent accessibility across the protein structure.

*Visualization in PyMOL:* The modified PDB file with updated B-factors was then loaded into PyMOL. The structure was colored according to the RSA values, where different colors represented varying levels of solvent exposure. For example, residues with high RSA values (indicating high solvent exposure) were colored differently from those with low RSA values (indicating low solvent exposure). Additionally, we used PyMOL to create zoomed-in views of specific regions of interest, such as the N-terminal domain and the core region.

### Molecular Dynamics Simulations

The initial nsp4 model, manually built based on the cryo-EM structure of the nsp4 chain from PDB entry 7Y38 to adopt an extended conformation consistent with the SAXS envelope, was subjected to all-atom molecular dynamics simulations using AMBER software suite (45) with parameters from the amber14SB forcefield (46).

All-atom MD simulations were performed in explicit water (TIP3P model (47)); a minimal distance of 20 Å between the edge of the solvent box and any protein atoms was imposed.

A cutoff radius of 12 Å was used in the calculations of non-bonded interactions with periodic boundary conditions applied; particle mesh Ewald method (48,49] was used to treat electrostatic interactions. To constrain hydrogens bonded to heavy atoms SHAKE algorithm {Ryckaert, 1977 #99)was employed. The simulations were performed by first relaxing the systems in two cycles of energy minimization; subsequently, the systems were slowly heated to 300 K using the parallel version PMEMD under NVT conditions (constant volume and temperature). Langevin dynamics (50) with collision frequency (γ=2) was used to regulate temperatures. The heated systems were then subjected to equilibration by running 100 ps of MD simulations under NPT conditions (constant pressure and temperature) with 1 fs integration time step. MD trajectories were collected over 200 ns at 1 ps interval and 2 fs integration time step. Analyses of the trajectories from MD simulations were done using CPPTRAJ program (51). MD simulations were carried out on a multi-GPU workstation with 2x AMD EPYC 7702 64-core processor and 2x Nvidia RTX A5000.

## RESULTS

### Purification of ONNV nsP4 RdRp and characterization of its RNA-binding and polymerase activities

The first alphavirus isolated was Western Equine Encephalitis Virus (WEEV) in the 1930’s (20). By the mid-1990’s, the complete genome sequence of Sindbis virus (SINV) was known, with reverse-genetic systems and replicons following in rapid succession (30,52–58). However, a highly active, recombinant alphavirus nsP4 RNA-dependent RNA polymerase is still not available, despite several published attempts (21–25). The earliest attempt was guided by alignment of the SINV nsP4 RdRp with those from enteroviruses (22). This study highlighted the presence of an ∼160 amino acid extension of the amino terminus of SINV and other alphavirus nsP4 RdRps (22). Since this region of the protein was devoid of sequences important for polymerase activity, the assumption was made that it might not be needed (22,23,25). Preparations of amino-terminally deleted derivatives of the SINV nsP4 RdRp only exhibited terminal transferase activity (22). Later, a full-length preparation of the SINV nsP4 RdRp was evaluated and shown to require the presence of other non-structural proteins to demonstrate any activity, although which non-structural proteins were required was not clear (21).

More recent studies of the alphavirus nsP4 RdRp have produced crystal structures for the truncated enzymes from SINV and Ross River virus (RRV) (23). This same study also pursued information on the full-length protein, but the major conclusion was that the amino-terminal domain is intrinsically disordered (23). While polymerase activity on a primed template formed by a hairpin RNA was reported, these data were later disproved by this group (23,25).

The most recent studies of a full-length alphavirus nsP4 RdRp used the sequence of that from O’nyong-nyong virus (ONNV) (23). The decision to use the ONNV nsP4 RdRp was made after evaluating the solubility and stability of full-length nsP4 proteins from many alphaviruses (23,24). To observe any activity required the presence of at least nsP1 and its ability to form a dodecameric ring, but the observed activity required hours for fractional conversion of the hairpin RNA to dsRNA (23). The requirement for the nsP1 dodecamer appeared to be to hold the nsP4 RdRp, as the nsP4 RdRp was in the central hole formed by the dodecameric ring of nsP1 molecules based on models built using cryogenic electron microscopy (cryo-EM) (23). While this study assumed that ONNV nsP4 adopted the same structure predicted by AlphaFold, there was no empirical evaluation of the solution behavior of the ONNV nsP4 RdRp as isolated from E. coli (23).

Since full-length ONNV RdRp was deemed the best-behaved alphavirus RdRp (24), we decided to purify and characterize this protein with the goal of inspiring hypotheses for the absence of polymerase activity. We expressed the ONNV nsP4 RdRp SUMO-tagged fusion protein to ensure the ability to produce the protein with an authentic Tyr amino terminus (**Supplementary Fig. S1A**) (33). A substantial fraction of the protein was soluble. We used polyethyleneimine (PEI) to precipitate nucleic acid followed by ammonium sulfate precipitation to remove free PEI (**Supplementary Fig. S1B**). These steps were critical to reproducible chromatographic behavior of the protein and removal of contaminants like the T7 RNA polymerase. A combination of Ni^2+^ affinity chromatography and ion exchange chromatographies was sufficient to purify the protein to greater than 95% purity (**Supplementary Fig. S1C-E**). The protein used for this study is shown in **Fig. 1**.

**Figure 1.**
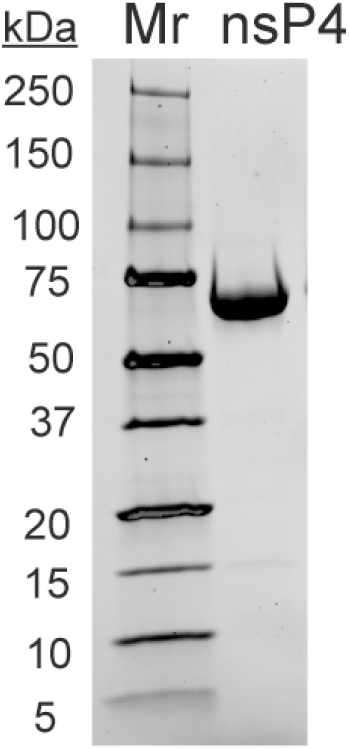
SDS-PAGE analysis of purified alphavirus nsP4 RdRp. SDS-PAGE analysis of bacterially expressed and purified nsP4 (69 kDa). Shown is a 4-15% polyacrylamide gel with a total of 5 µg of nsP4. Broad-range molecular weight markers (Mr) and corresponding molecular weights are indicated. Refer to full purification in Supplementary Fig. S1.

We evaluated the ability of the protein to bind to ssRNA 19 (rU_10_ss9) or 25 (rC_25_) nucleotides in length by using fluorescence polarization (see Materials and Methods). ONNV RdRp exhibited an affinity for RNA in the 4 -15 µM range (Alpha RdRp in **Fig. 2A-B**). In contrast, the RdRps from poliovirus (Entero) and Zika virus (Flavi) exhibited much higher affinity for ssRNA (**Fig. 2A-B**).

**Figure 2.**
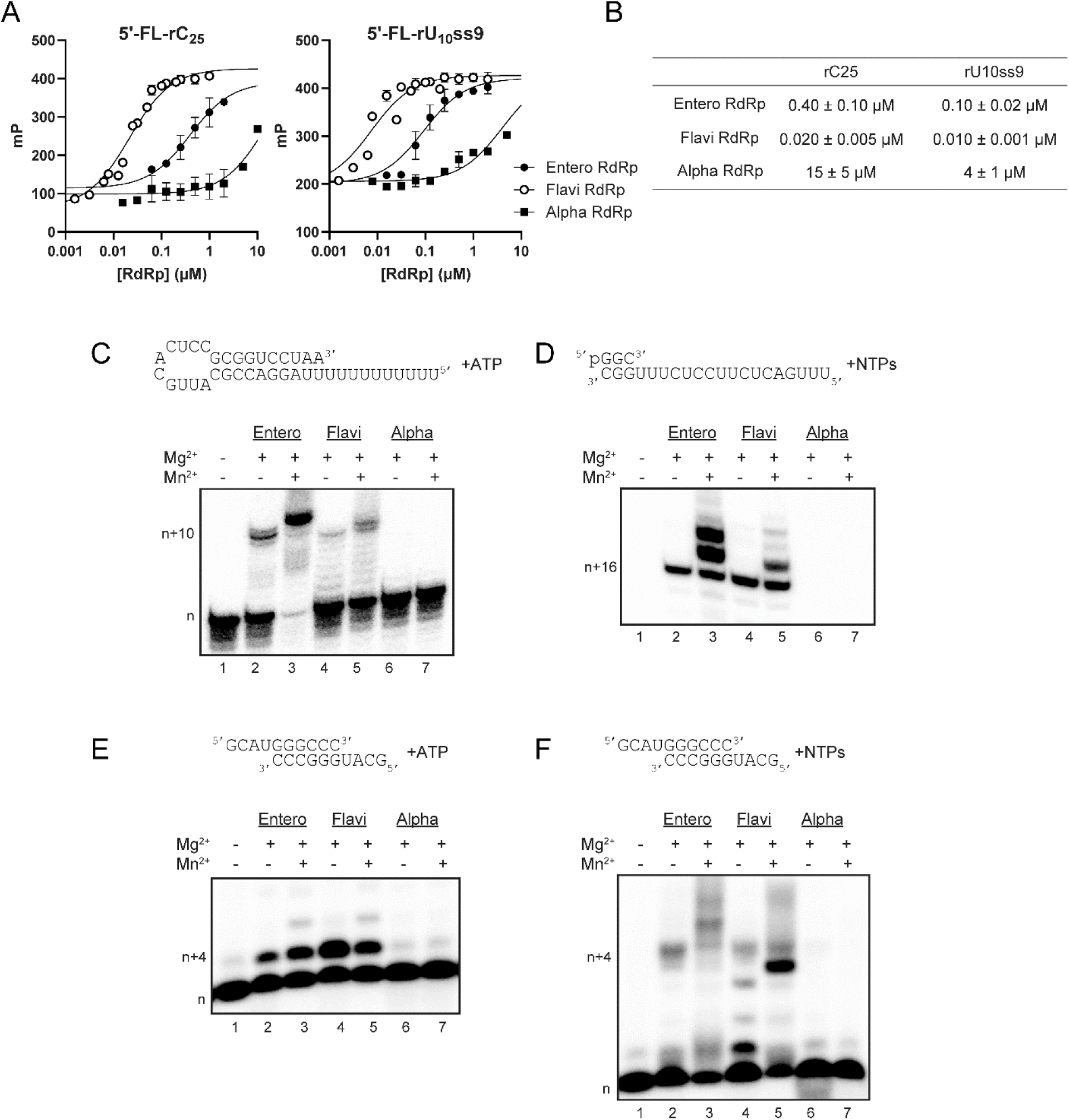
Alphavirus nsP4 RdRp binds RNA with low affinity and does not exhibit RdRp activity. (**A,B**) RNA binding assays. Fluorescein-labeled ssRNA (rC25 or rU10ss9) was mixed with increasing concentrations of either an Enterovirus RdRp, Flavivirus RdRp, or Alphavirus RdRp. Millipolarization (mP) was plotted and the data were fit to a hyperbola, yielding an apparent dissociation constant as reported in the table in panel B. The Alphavirus RdRp binds RNA with an affinity orders of magnitude lower than both the Enterovirus and Flavivirus RdRp. (**C-F**) RdRp activity assays. Primer extension assays utilizing three different RNA primer-template substrates in the absence and presence of Mn2+. We utilized a short RNA hairpin substrate (panel C), trinucleotide primer with 20-mer RNA template (panel D), or a symmetrical primer-template substrate (panel E & F). Reactions contained RNA substrate, corresponding ATP or NTPs and RdRp, were incubated at 30 °C for 30 min and quenched. Reaction products were resolved by denaturing PAGE and visualized by phosphorimaging. The Enterovirus and Flavivirus RdRps exhibit primer-extension activity, however, the Alphavirus RdRp does not. The presence of Mn decreases the specificity of nucleotide incorporation for both the Enterovirus and Flavivirus resulting in non-template nucleotide addition and greater product formation.

We evaluated the polymerase activity using a variety of primed templates, including the following: a hairpin template reported for alphavirus RdRps (**Fig. 2C**) (33); a trinucleotide-primed template reported for flavivirus RdRps (**Fig. 2D**) (35,59); and the symmetrical primed-template reported for enterovirus RdRps (**Fig. 2E**) (60). ONNV RdRp failed to use any of these substrates while the entero and flavi RdRps used all three substrates (**Fig. 2C-E**). We attempted to relax the specificity of the reaction by adding Mn^2+^ to the reaction (24,61,62). While this approach clearly worked for the entero RdRp (compare lane 3 to lane 2 in **Fig. 2C**), there was no impact on the outcome for the alpha RdRp (compare lane 7 to lane 6 in **Fig. 2C**). Similar observations were made for the other substrates as well (**Fig. 2D-E**).

### Biophysical characterization of ONNV nsP4 RdRp

#### Circular dichroism spectroscopy (CD)

The initial characterization of the structure and dynamics of SINV and RRV nsP4 RdRps suggested an intrinsically disordered and conformationally flexible amino-terminal domain (23). However, the AlphaFold model of ONNV nsP4 revealed a folded, stable conformation for this domain (24). We used this model to calculate the helical content and found 35% α-helix and 12% β-sheet. We reasoned that empirical analysis of secondary structure of ONNV nsP4 by using circular dichroism spectroscopy would distinguish between the two reported possibilities. This empirical analysis was consistent with 33% α-helix and 18% β-sheet (**Fig. 3**).

**Figure 3.**
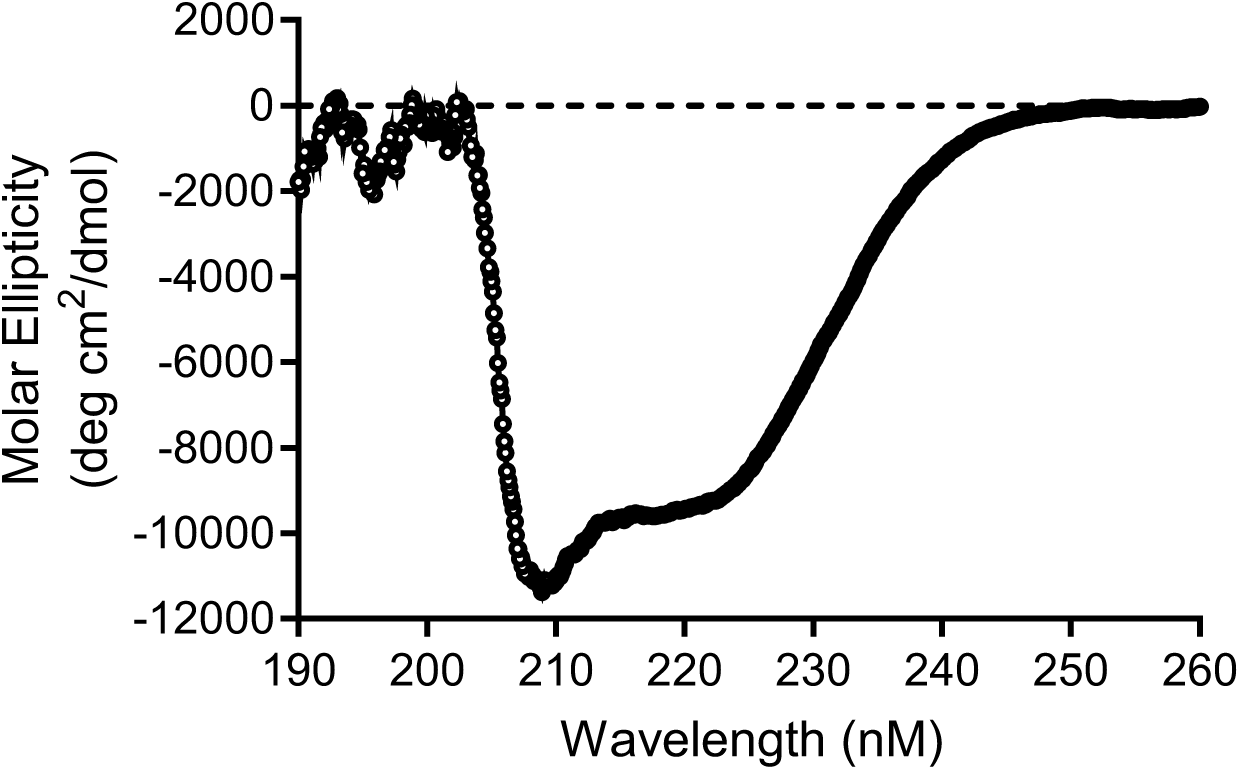
Circular Dichroism spectrum of nsP4 was consistent with predicted helical and β-sheet content. BestSel method for protein secondary structure prediction as from the CD data indicates 33.1% helix and 18% sheet. The calculated CD secondary structure map as from Pdb2CD program with the nsP4 cryo-EM structure points to 35% helix and 12% sheet, in agreement with the experimental values and suggesting that the NTD perhaps retains all of its secondary structure in solution.

*Size-exclusion chromatography and multi-angle light scattering (SEC-MALS)* – It has been noted that full-length alphavirus polymerases are prone to aggregation in solution (23).

Size-exclusion chromatography fractionates based on the hydrodynamic radius, so it can only be used to give a relative molecular mass. When coupled to the measurement of refractive index (RI) and UV detectors, both of which give an accurate protein concentration, and multi-angle and dynamic light scattering detectors, an absolute molecular weight can be determined (63). We evaluated our ONNV nsP4 preparation using SEC-MALS-DLS. The data were consistent with a species of 69.7 kDa compared to a calculated molecular weight of 68.3 kDa (**Fig. 4**).

**Figure 4.**
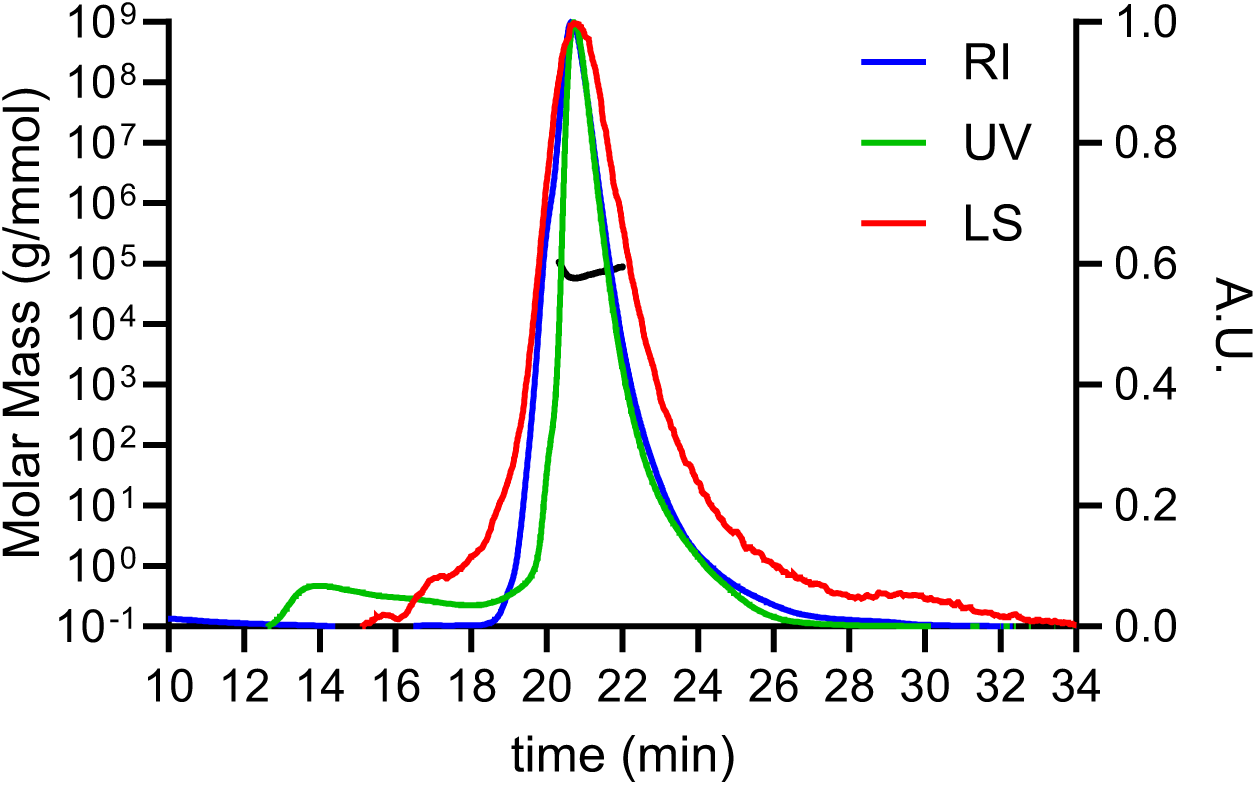
SEC-MALS of nsP4 is consistent with a single species with a MW of 69.7 kDa. Analysis indicates a chromatogram with a single peak in the light scattering (red), UV (green) and the refractive index (blue) detectors. The main peak corresponds to a monodisperse species reflecting a monomer of nsP4 with a molar mass of 69.7kD.

#### Analytical ultracentrifugation (AUC)

Analyses to this point confirmed a well behaved, structurally ordered monomer in solution, making the protein a candidate for AUC. AUC offers the ability to determine the sedimentation coefficient, the molecular weight, and the frictional ratio. The sedimentation coefficient measures the rate of sedimentation over time. If conformational changes occur during the sedimentation experiment, then the sedimentation rate will change over time, producing an asymmetric curve. In addition, we can obtain information on the shape of the protein. For a 60-80 kDa protein, a globular protein will sediment at 4-6 S while an elongated protein will sediment at 2-3 S. A second measure of shape is the frictional ratio. For globular proteins the value would be in 1.1 – 1.3 range while an elongated molecule would exhibit a value greater than 1.8.

For ONNV, the sedimentation rate appeared constant over time with a coefficient of 3.2 S (**Fig. 5A**), corresponding to a molecular weight of 69 kDa (**Fig. 5B**). The frictional coefficient was also consistent with a single sedimenting species; the value was 1.8 (**Fig. 5C**). Finally, greater than 80% of the signal measured was attributable to a single 69 kDa species with a frictional coefficient of 1.8 (**Fig. 5D**). These data suggest that ONNV nsP4 exists in an elongated conformation in solution instead of the globular conformation predicted by AlphaFold (24).

**Figure 5.**
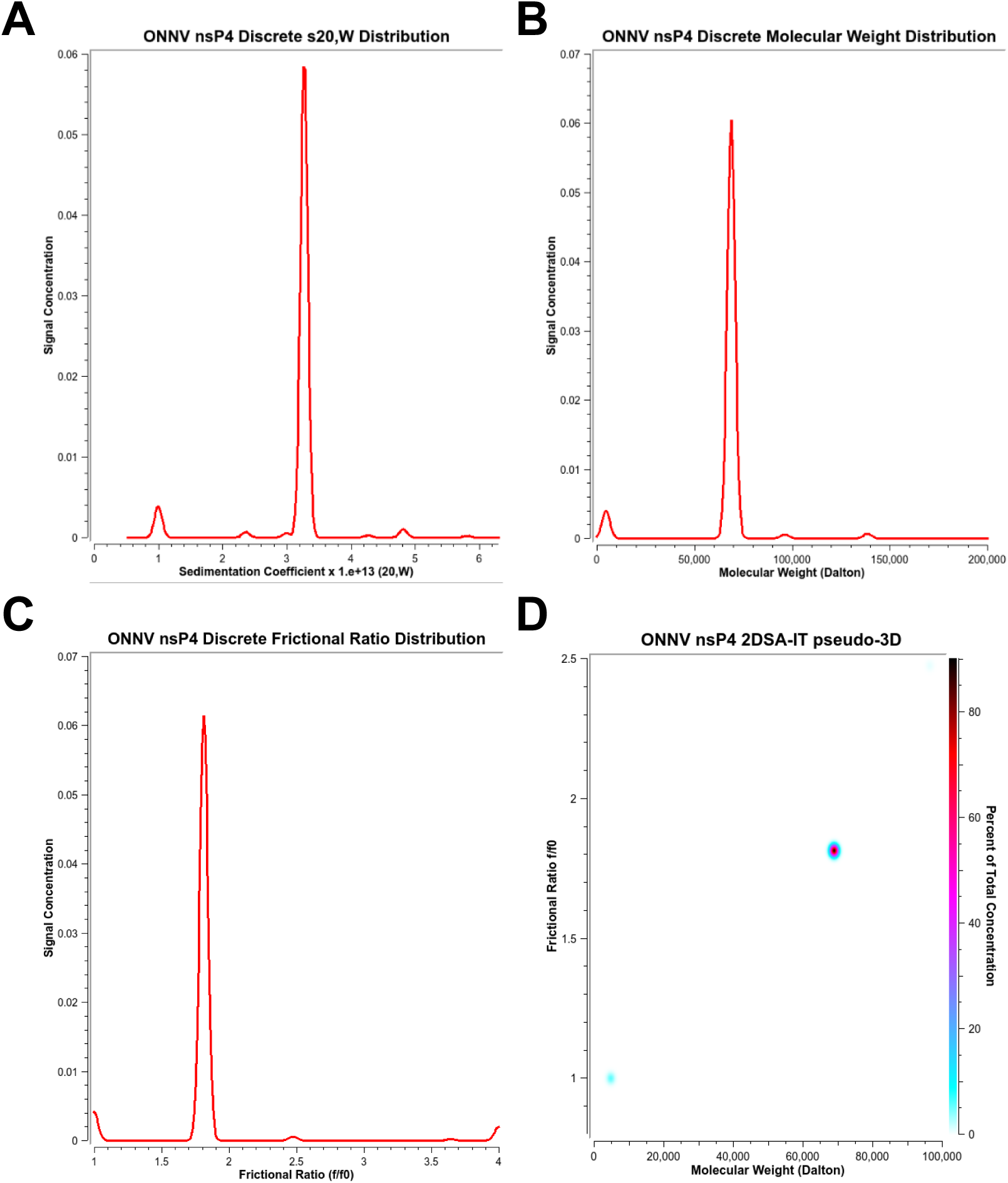
Analytical ultracentrifugation of nsP4 indicates a highly homogenous, single-species with an elongated conformation. (**A**) The S,20w distribution of nsP4 shows a stable sedimentation coefficient of 3.2S. (**B**) Molecular weight distribution of the same AUC data, showing an estimated molecular weight of 69 kDa, which is in excellent agreement with the estimated monomeric MW = 68,380. (**C**) Frictional coefficient ratio (f/f0) distribution, showing a single value of 1.8 for the monomer with no spread, suggesting a highly elongated protein with no large disorder (in agreement with SAXS). (**D**) Pseudo-3-dimensional plot of the AUC data, the peak for the monomer is >80% of the signal observed, suggesting a highly homogeneous and conformationally ordered sample.

#### Small-angle X-ray scattering (SAXS)

SAXS offers the opportunity to define the overall shape of a protein in solution. The raw scattering data was smooth and continuous, indicating a dataset of high quality (**Supplementary Fig. S2**). The Guinear plot was linear, consistent with the absence of aggregation and the absence of interparticle interference (**Supplementary Fig. S3**) (64). The Kratky plot was bell shaped and returned to baseline after reaching a peak, as expected for a well-folded protein lacking substantial disorder (**Supplementary Fig. S4**) (64). Finally, the pair-distance distribution was smooth and lacking any unnatural oscillations (**Supplementary Fig. S5**) (64). The longest pairwise distance calculated from the distribution was 107 Å, again consistent with an elongated molecule.

Given the high-quality SAXS data obtained, we used the DENSS (DENsity from Solution Scattering) algorithm to calculate the molecular envelope of for ONNV nsP4 in solution (**Fig. 6**) (64). It was immediately evident that the AlphaFold model of ONNV nsP4 would not fit into the calculated envelope (**Fig. 6A**). A rigid body rotation of the amino-terminal domain in an unstructured “hinge” region of the protein centered around Pro-103 revealed a conformation that fit the observed electron density well (**Fig. 6B**). Consistent with this observation was the finding that the theoretical scattering produced using the AlphaFold model yielded a poor fit when compared to the empirical dataset based on the Chi-square value of 7.8 for the Alphafold model versus 2.6 for the elongated model (**Supplementary Fig. S6**) (64). A good fit would have a Chi-square value in the 1.0 to 2.0 range.

**Figure 6.**
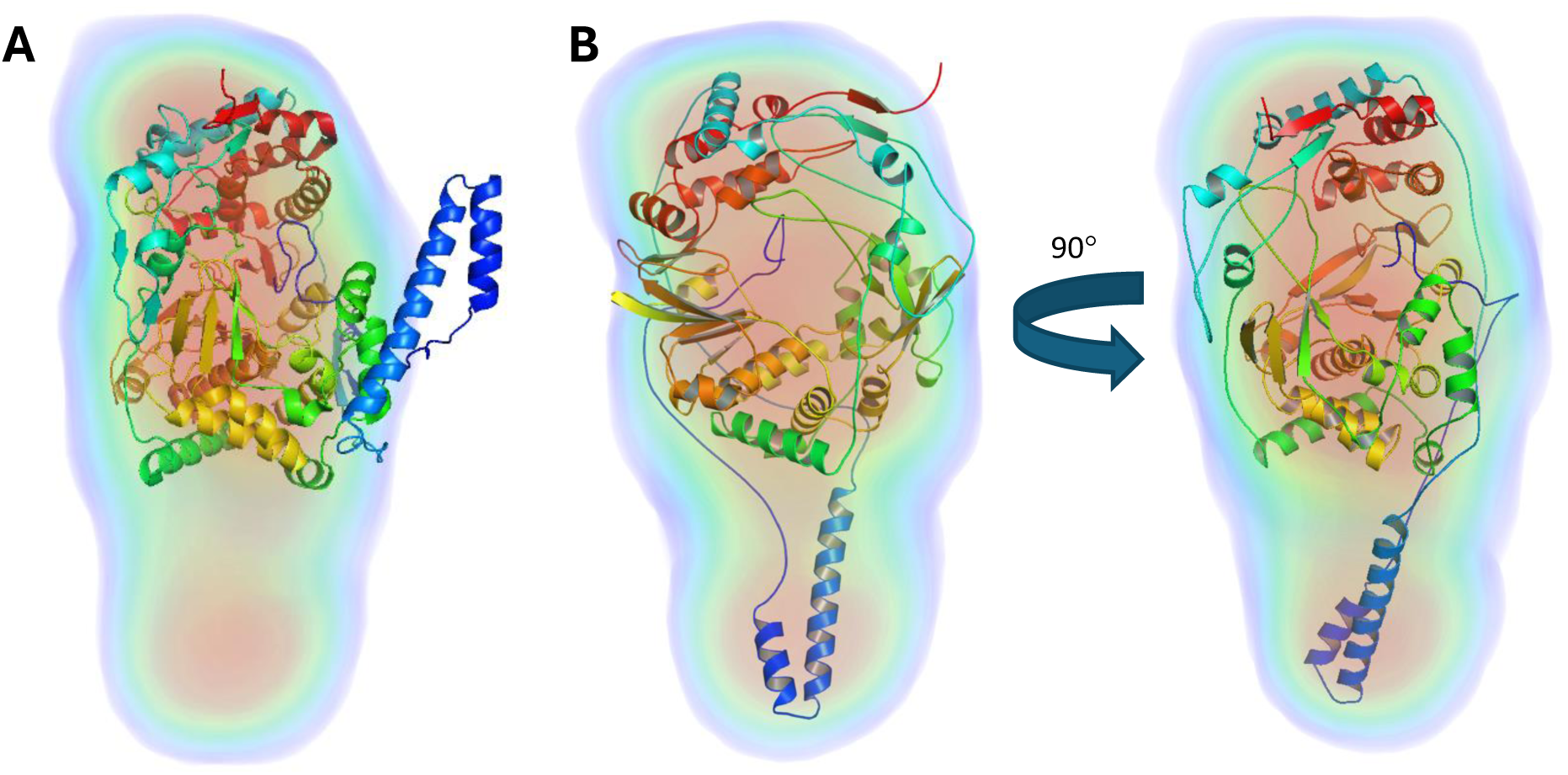
SAXS of nsP4 displays an elongated shape that requires a rotation of the N-terminal domain for proper fitting. (**A**) Placing the compact structure of nsP4 in the SAXS envelope with no adjustments shows that this structure does not fit the electron density. (**B**) Two perpendicular views of the DENSS-calculated envelope electron density were fitted to the nsP4 model. SAXS-derived fits as from replicates done at two concentrations, 2.9 mg/mL and 1 mg/mL both indicated an extended conformation for the N-terminal domain of nsP4. This needed a 180-degree flip of the 100-residue N-terminal region, resulting in an alternative open and inactive conformation of nsP4.

#### Hydrogen-deuterium exchange mass spectrometry (HDXMS)

We have used HDXMS as an orthogonal approach to probe the conformational ensemble behavior of ONNV nsP4 in solution and to determine the extent to which any conformational dynamics of the protein can be observed. HDXMS is ideally suited for assessing the conformational state of ONNV nsP4 because deuterium exchange provides a readout of hydrogen bonding in a protein structure(65). Additionally, slow interconversion between conformations can be observed by the presence of bimodal hydrogen-deuterium exchange profiles reflective of the EX1 exchange regime (66,67). The deuterium-exchange reaction was performed over a 1 – 100 min time period. We detected 184 peptides spanning 91.8% of the ONNV nsP4 sequence (**Supplementary Fig. S7**).A back exchange correction was also applied using a maximally deuterated control (37) in Deuteros 2.0 (38). Only EX2 kinetics were observed for all regions except for one carboxy-terminal locus (residues 560-580), consistent with the observation of only one conformation in solution at the seconds-minutes deuterium exchange experimental timescales and consistent with AUC and SAXS (**Supplementary Fig. S8**).

We mapped relative fractional deuterium uptake onto both the compact and extended conformations to determine which conformation was more consistent with the observed deuterium exchange (**Supplementary Fig. S9**). Regions with common secondary structure between the extended and compact conformations had deuterium exchange profiles consistent with their structures (**Supplementary Fig. S10**). Two loci were identified with near amino acid resolution that had a different conformation in the extended structure (**Fig. 7A,B**). In the compact structure, residues 26-29 form a β-sheet with residues 98-101, while in the extended structure, both of these regions are unstructured and have no specific contacts. After 1 min of deuterium exchange, peptide 23-28 exhibits near maximal deuterium exchange (**Fig. 7C**), consistent with the extended structure with low H-bonding propensities (**Fig. 7A,C**). In the extended structure, residues 30-33 form contacts with residues 234-236, while both regions exist as unstructured loops in the compact conformation (**Fig. 7B**). Peptide 228-237 shows low to intermediate deuterium uptake while peptide 29-40 shows one amide that remains protected even after 100 min deuterium exchange (**Fig. 7B,D**), consistent with the extended structure (Englander and Kallenbach 1983) https://pubmed.ncbi.nlm.nih.gov/6204354/.

**Figure 7.**
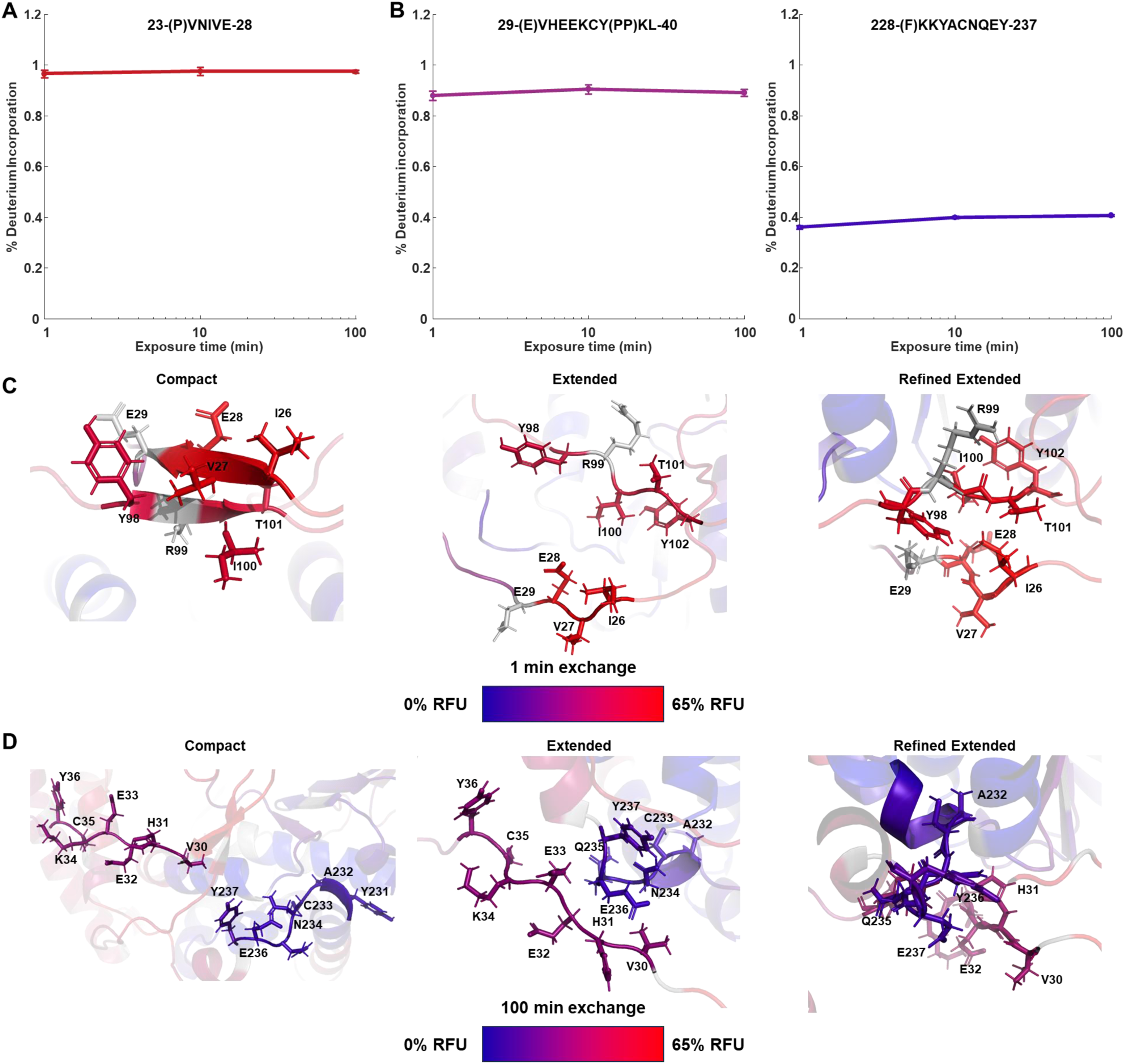
Deuterium uptake at N-terminal loci is consistent with the extended conformation of nsP4. Deuterium uptake plots for **(A)** peptide 23-28 and **(B)** peptides 29-40 (left) and 228-237 (right) plotting % deuterium incorporation corrected for % deuterium in labelling buffer and back exchange after 1, 10, and 100 min on a semi-log scale. Average back exchange estimates were used for peptide 228-237 which was not observed in maximum deuteration experiments. (**C**) Relative fractional deuterium uptake mapped onto N-terminal residues 26-29 and 98-101 of the compact (left), extended (middle), and refined extended structures of nsP4 after 1 min exchange. Uptake is displayed as a blue (low deuterium uptake) to red (high deuterium uptake) gradient. (**D**) Relative fractional deuterium uptake mapped onto residues 30-36 and 232-237 of the compact (left), extended (middle), and refined extended structures of nsP4 after 100 min exchange. Uptake is displayed as a blue (low deuterium uptake) to red (high deuterium uptake) gradient.

### Mutually exclusive conformations of the ONNV RdRp

Our biophysical experiments were inspired and evaluated initially by using the model created manually (see accompanying PDB file) based on PDB 7Y38. Given the overwhelming congruence of the biophysical data supporting an extended conformation, we used molecular dynamics simulations to refine the model of the extended conformation (see Materials and Methods and accompanying PDB file). The final model is shown in **Fig. 8A**. This model was accommodated well by the SAXS envelope (**Fig. 6B**) and further strengthened our interpretation of the HDXMS experiments (**Fig. 7A,B** and **Supplementary Figs. S9,S10**).

**Figure 8.**
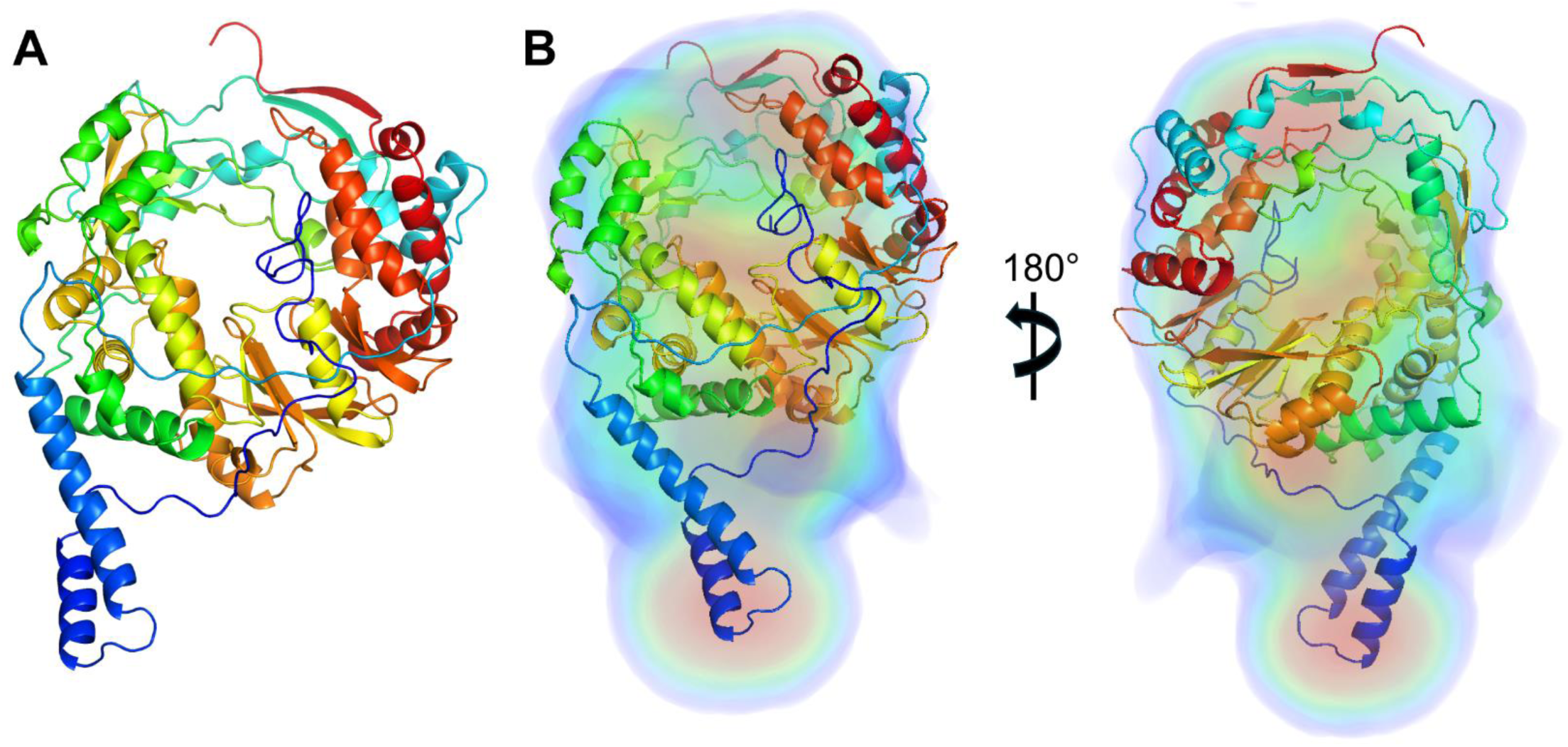
Refined model of the extended conformation fits the SAXS envelope. (**A**) Molecular dynamics simulation of extended conformation. (**B**) Placing the structure of nsP4 in the SAXS envelope.

The most striking observation was the apparent existence of two, mutually exclusive conformations of the first 100 amino-terminal residues. The helix bundle (residues 43 – 83) adopted distinct orientations in the extended and compact conformations (compare the pink helices in **Fig. 9A** to those in **Fig. 9B**). The distinct conformations of residues flanking the helices, particularly residues 20 – 38 and 80 – 90, were even more intriguing as very clear differences in predicted interactions could be identified (compare red residues in **Fig. 9A** to those in **Fig. 9B**; summarized in **Table 1**).

**Figure 9.**
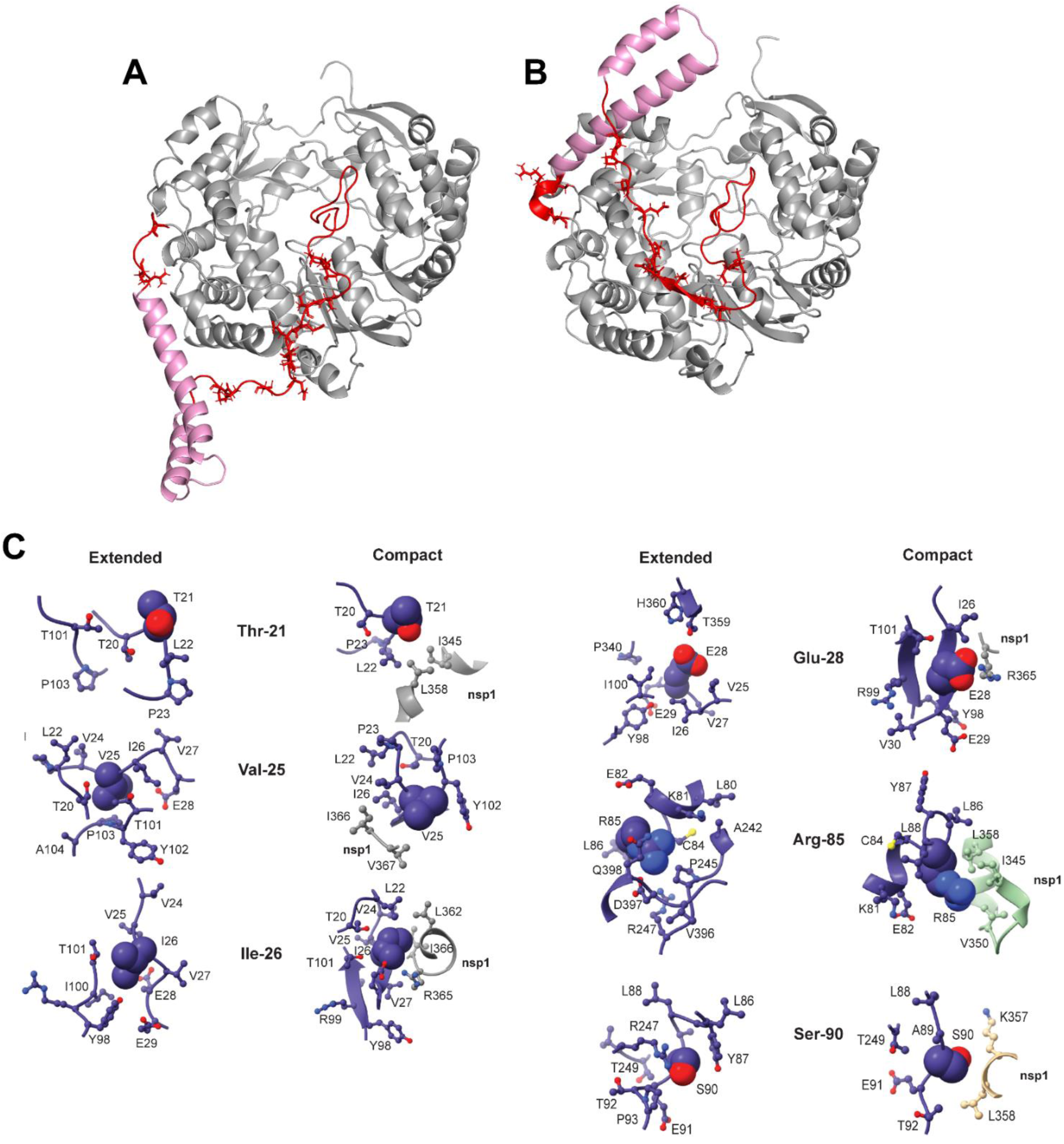
Comparison of the N-terminal interactions in the extended and compact conformations reveal distinct interactions between the two states. (**A**) Extensive interactions between the N-terminal domain (aa 1-90) and the rest of nsP4 polymerase in the extended conformation. Red: aa 1-42 and 84-90; Pink: aa 43-83. Sidechains that are not in NTD helices with interactions as shown in Table 1 are shown as sticks. (**B**) Extensive interactions between the N-terminal domain (aa 1-90) and the rest of nsP4 polymerase in the compact conformation. red: aa 1-42 and 84-90; pink: aa 43-83. Sidechains that are not in NTD helices with interactions as shown in Table 1 are shown as sticks. (**C**) Close-up view of interactions of residues in the NTD that change from the compact to extended conformation.

**Table 1.**
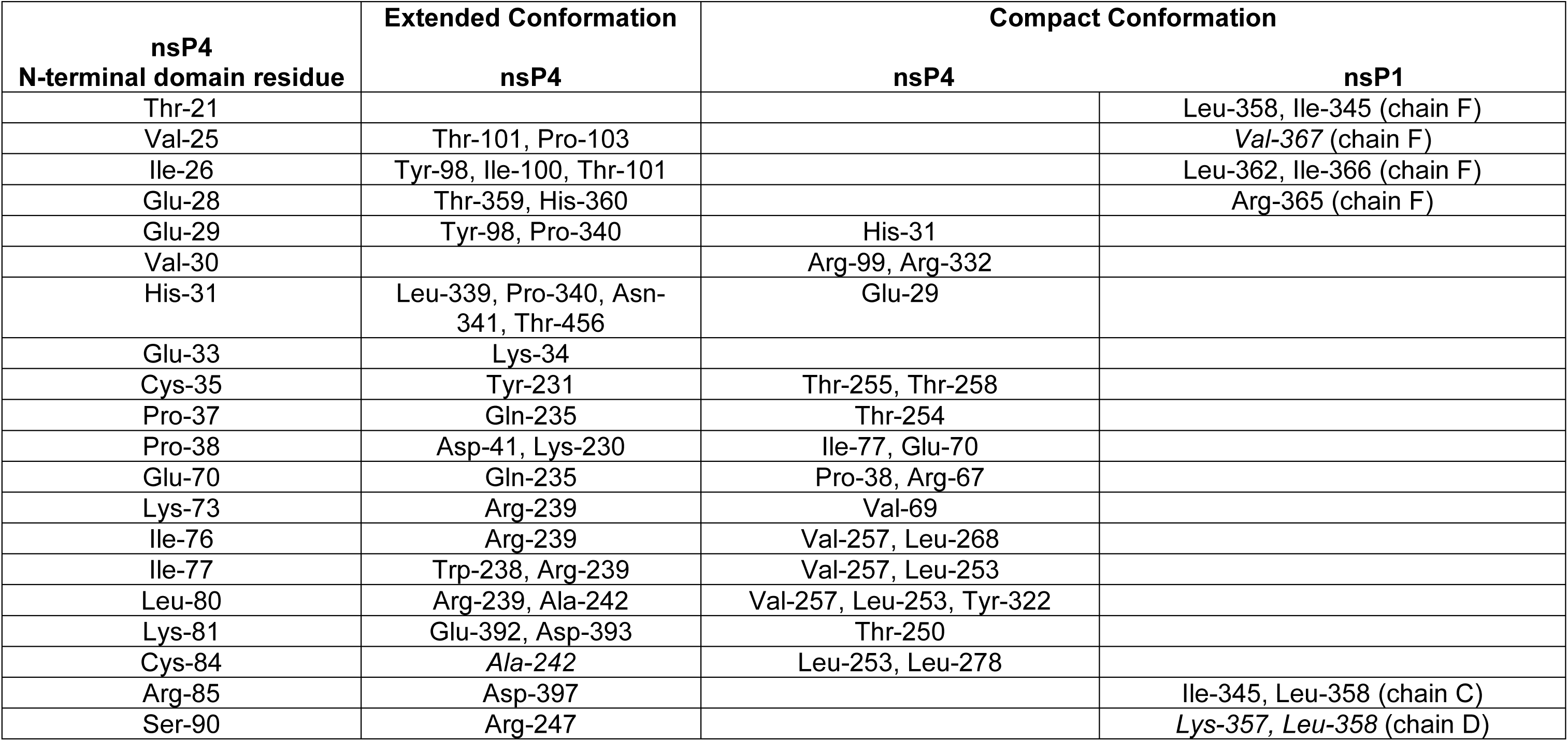
Mutually exclusive interactions between the N-terminal domain of ONNV nsP4 and the nsP4 core or nsP1 in the extended and compact conformations. The model of the extended form of ONNV nsP4 after molecular dynamics simulation was compared to the published model of the compact form of ONNV nsP4 that was restrained using the structure predicted by AlphaFold. Residues of the amino-terminal domain unique to alphavirus polymerases form distinct interactions in the extended and compact conformations. Residues of nsP4 that interact with nsP1 in the compact conformation are sequestered by residues of nsP4 in the extended conformation. nsP1 forms a dodecamer. Each subunit has been labeled chain A – L; the specific chain used for interaction with nsP4 is indicated explicitly. The interacting residues listed represent sidechain-sidechain interactions. Residues listed in italicized type contribute to either backbone-backbone or sidechain-backbone interactions.

The conformation of nsP4 in complex with the nsP1 dodecamer has amino-terminal residues Thr-21, Val-25, Ile-26, Glu-28, Arg-85, and Ser-90 interacting with residues of nsP1 from three different polypeptide chains (see Compact Conformation in **Table 1**). These very same residues are predicted to interact with the core of nsP4 when in the extended conformation (**Table 1**). The residues in between the nsP1-interacting residues also exhibit conformation-specific interactions (**Table 1**).

The nsP1-interacting residues may contribute to the nsP1-dependence for activity. These same residues may drive formation of the extended, inactive conformation. We compared each of these residues in the two different conformations. In the extended conformation, the sidechains of all nsP1-interacting residues of nsP4 exhibit extensive interactions with other nsP4 residues (see extended conformation panel in **Fig. 9C**). The extended conformation would therefore lack the ability to interact with the spherule in a manner similar to that of the compact conformation. However, these residues would likely not drive the conformational switch as there appear to be the same number of interactions between nsP4 and nsP1 in the compact state as between nsP4 residues in the extended state (compare compact conformation to extended conformation in **Fig. 9C**).

### Evolutionary conservation of both the extended and compact conformations of ONNV RdRp

Our analysis has emphasized ONNV nsP4. The final question that we addressed was: can both the extended and compact conformations exist in *all* alphavirus nsP4 RdRps? To address this question, we performed a phylogenetic analysis as described under Materials and Methods. We used the most diverse set of sequences available for alphaviruses to produce an alignment (**Supplementary Fig. S11**). First, we used the sequence alignment to quantify sequence conservation. We define the conservation using the effective number (Neff). Values range from 1.00 (most conserved) to 20.00 (least conserved) (**Supplementary Fig. S11**). A value of 1.00 means that only one amino acid is found at that position in the alignments while a value of 20.00 means that all 20 amino acids can be found at that position. For nsP4, the range was from 1.00 to 6.00, suggesting strong conservation of this protein across the alphavirus genus. We also used these alignments to perform a residue co-variation analysis (direct-coupling analysis, DCA (68)) with the expectation that some amino-terminal residues that are not highly conserved would have unique, co-varying residues in each conformation. The DCA is presented in **Supplementary Tables S1A,B** with pairwise distances mapped on the extended or compact conformations, respectively. To perform this analysis with the least bias possible, we filtered the DCA data to identify residues with the highest values for the Z-score, which indicates residue pairs with the highest probability of functional relevance (**Supplementary Fig. S11**). Residue pairs with a Z-score higher than 2.0 were deemed reliable for our analysis. Z-scores with values less than 1.00 generally mean that the residue pair is not co-varying, often because of the high conservation of one or both residues. Highly conserved pairs were also deemed reliable.

We focused first on residues implicated in unique interactions in the extended and compact conformations as presented in **Table 1** (**Table 2**). These residues exhibited conservation ranging from essentially absolute (1.00-2.00) to high (2.00-4.00) (**Table 2**). We observed a few very clear examples of co-variation of amino-terminal residues with residues in the extended and compact conformations. For example, Glu-29 co-varied with Tyr-98 in the extended conformation (Z-score of 4.32) and His-31 in the compact conformation (Z-score of 2.42). His-31 co-varied with Thr-456 in the extended conformation (Z-score of 2.41) and Glu-29 in the compact conformation (Z-score of 2.42). Leu-80 co-varied with Ala-242 in extended conformation (Z-score of 3.63) and Leu-253 in the compact conformation (Z-score of 2.96). These observations are consistent with evolutionary conservation of both conformations.

**Table 2.**
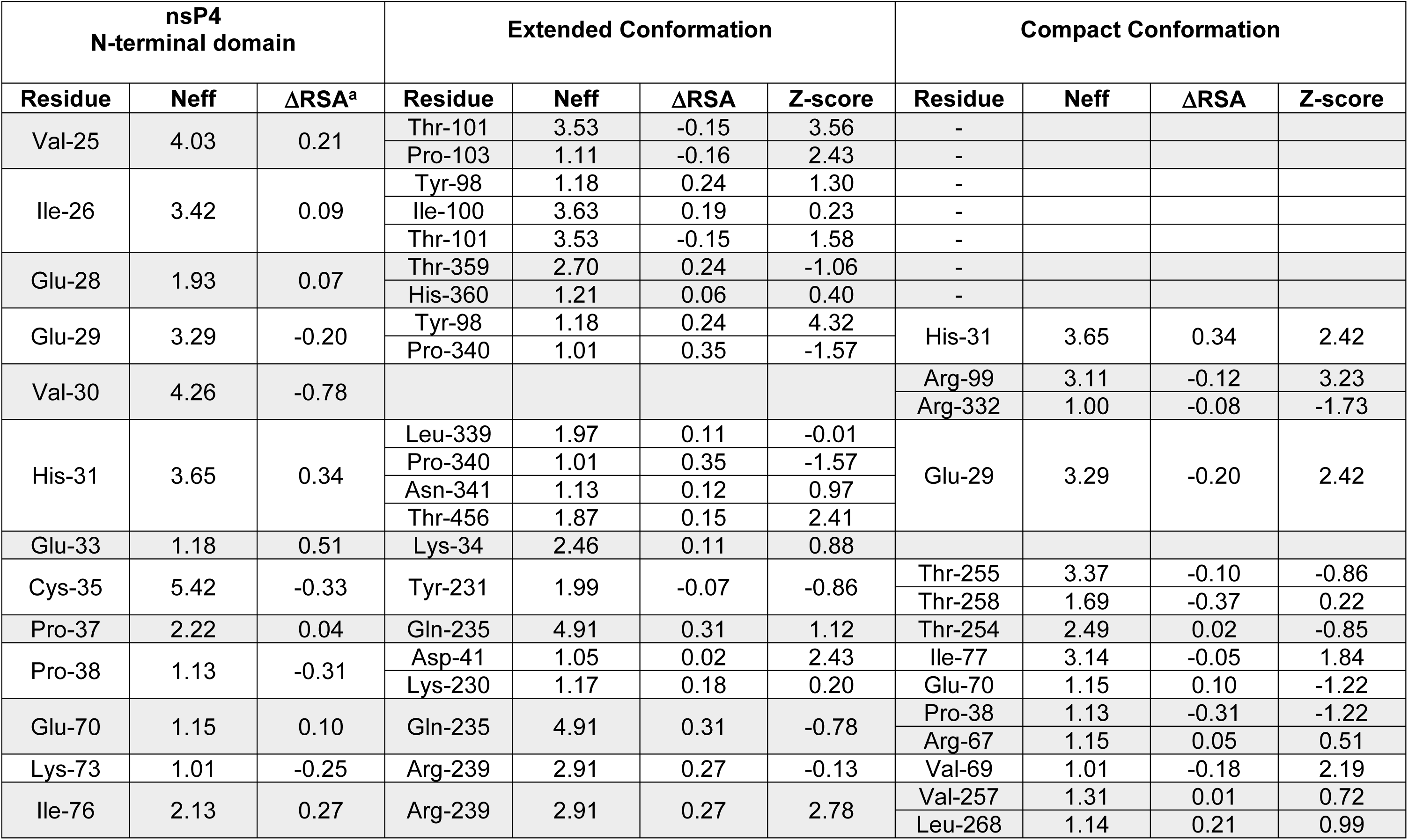
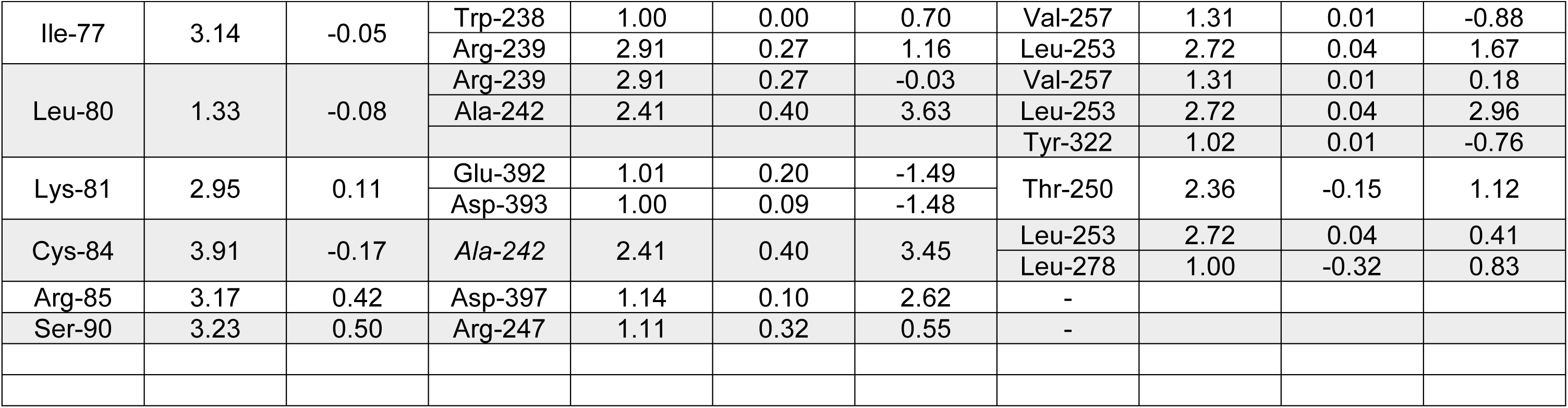
Residues implicated in unique interactions in the extended and compact conformations. Residues evaluated are from Table 1. When interacting residues are not indicated under *Compact Conformation*, the residue interacts with an nsP1 subunit. Neff is a measure of conservation on a scale from 1.00 (absolutely conserved) to 20.00 (no conservation). ΔRSA is the change in relative solvent accessibility. The range observed was −0.78 to 0.51. These values are expressed for the transition from extended to compact. A positive number indicates more accessible in the compact than in the extended. Z-score is a measure of the reliability of the indicated pairwise interaction based on co-variation analysis. Values greater 2.00 are reliable. Very low Z-scores are generally caused by the absence of any observed co-variation because residues are highly conserved.

We also used a more unbiased approach to select residues for interrogation, including investigation of any corresponding pairwise interactions. We evaluated the change in relative surface area of each residue of the amino terminus in going from the extended conformation to the compact conformation (see **Supplementary Table. S2**). The absolute value of the change in relative solvent accessibility (RSA) is shown in **Fig. 10A**. We evaluated the top five residues exhibiting the largest negative change (more accessible in extended than in compact; see residues in **cyan** in **Fig. 10B**) and the largest positive change (more accessible in the compact than in the extended; see residues in **black** in **Fig. 10B**). The values for the ΔRSA and Neff of these residues are provided in **Table 3**.

**Figure 10.**
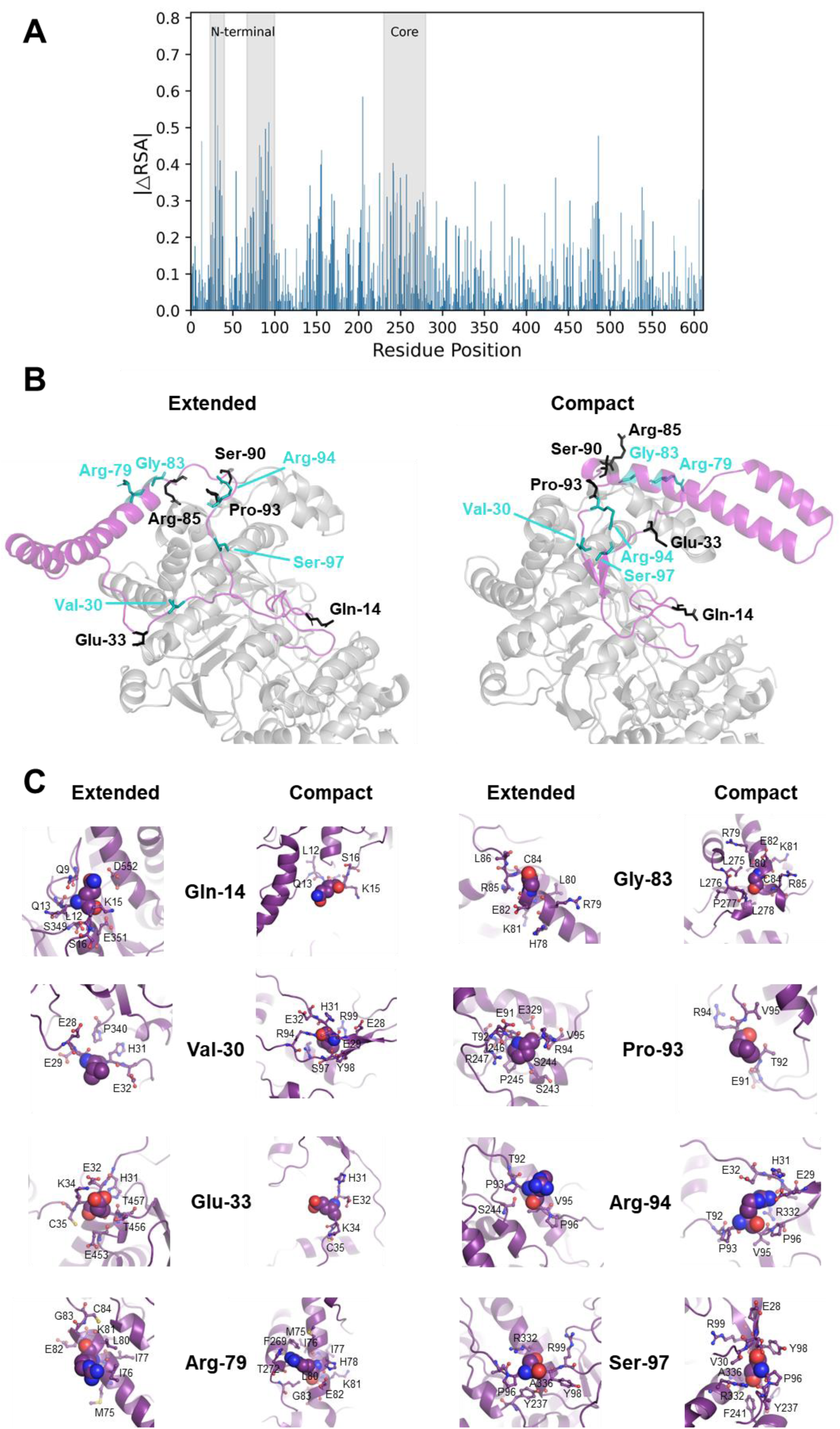
**Absolute differences in Relative Solvent Accessibility (RSA) in nsP4 and interrogation of residues in pairwise interactions suggest the extended and compact states being evolutionary conserved**. (**A**) Absolute differences in RSA between the two conformations (compact and extended) of the nsP4 protein, highlighting in gray the regions that undergo the most significant structural changes. Notably, two regions in the N-terminal domain (20-40 and 60-100) and one in the core region (residues 230–270), exhibit the largest changes in RSA, indicating a substantial residue relative solvent accessibility change and a potential interaction between the two regions. (**B**) Residues that exhibit the largest changes in rASA between the extended and compact conformation. Residues 1 to 100 are colored in purple. Residues with the largest change in rASA (see Table 3) are show as stick. The top 5 residues with the largest change in rASA in going from extended to compact with high to low accessibility are shown in teal and those with low to high accessibility are shown as black. (**C**) Close-up view of interactions of residues in the NTD that hve the largest change in ΔrASA between the extended and compact conformation (Table 3). The indicated residue is shown as CPK. Residues around the indicated residue are indicated.

**Table 3.**
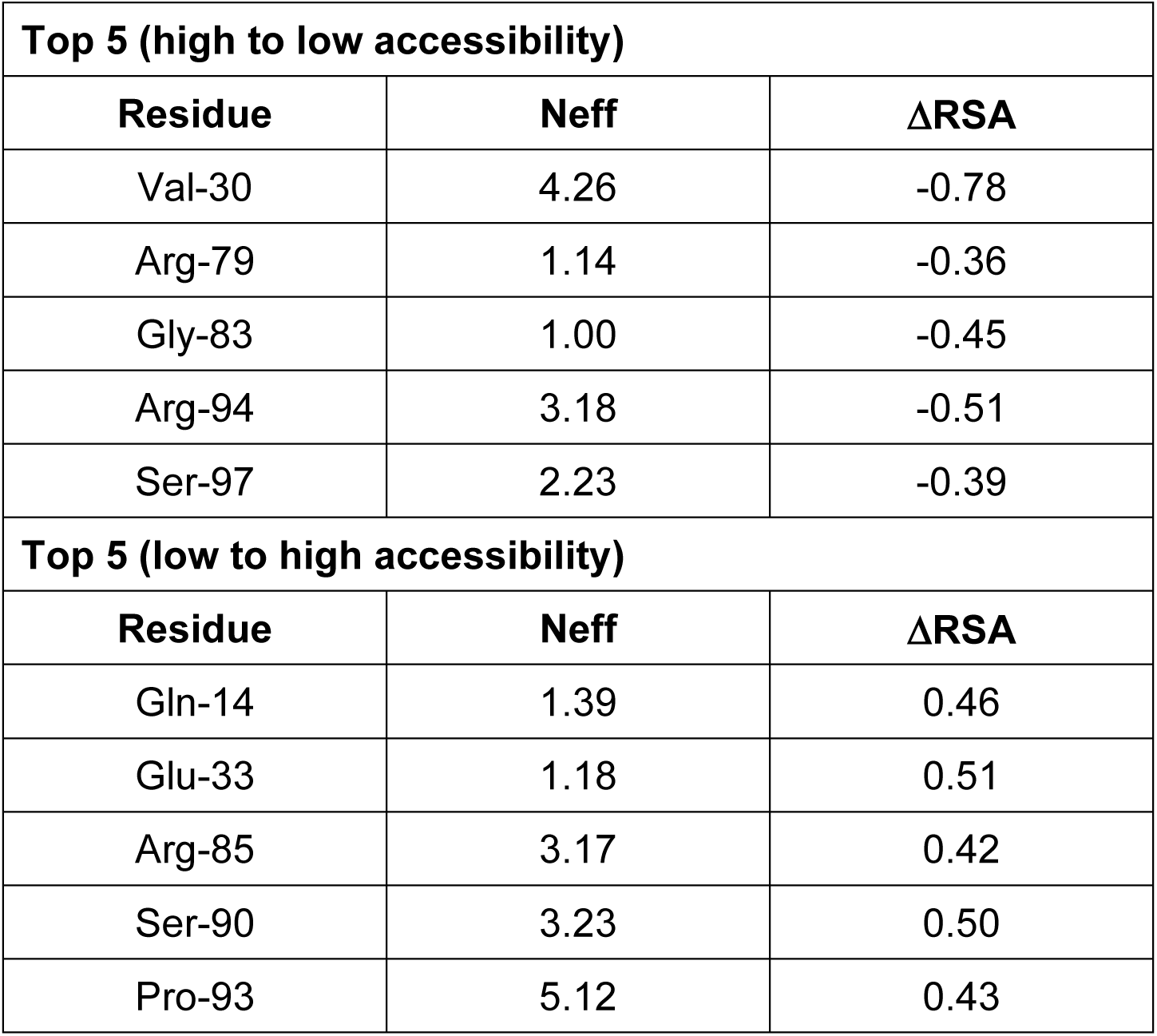
Residues of the N-terminal domain exhibiting the largest changes in ΔRSA. Shown are the top 5 residues with the largest values for ΔRSA in going from extended to compact (high to low accessibility) or in going from extended to compact (low to high accessibility). The order is sorted by amino acid residue number.

| Top 5 (high to low accessibility) |  |  |
| --- | --- | --- |
| Residue | Neff | ΔRSA |
| Val-30 | 4.26 | -0.78 |
| Arg-79 | 1.14 | -0.36 |
| Gly-83 | 1.00 | -0.45 |
| Arg-94 | 3.18 | -0.51 |
| Ser-97 | 2.23 | -0.39 |
| Top 5 (low to high accessibility) |  |  |
| Residue | Neff | ΔRSA |
| Gln-14 | 1.39 | 0.46 |
| Glu-33 | 1.18 | 0.51 |
| Arg-85 | 3.17 | 0.42 |
| Ser-90 | 3.23 | 0.50 |
| Pro-93 | 5.12 | 0.43 |

The most substantial changes occurred at residues 14-33 and 79-97, both of which are at the base of the loop that changes conformation in going from the extended state to the compact state (**Fig. 10B**). These residues ranged from highly conserved to moderately variable (compare values for Neff in **Table 3**). We observed three classes of changes: (1) those involving nsP1 in the compact state, Arg-85 and Ser-90 (**Fig. 9B**); (2) those involving distinct interactions in the extended and compact states, Gln-14, Gly-83, and Ser-97 (**Fig. 10C**); and (3) those that exhibit interactions in only one conformational state, Val-30, Glu-33, Arg-179, Pro-93, and Arg-94 (**Fig. 10C**).

We recognize that the structural model of the extended state is a model that will require further experimental investigation and/or empirical structure determination. However, we would be remiss to ignore the residues that have distinct interaction in both the compact and extended conformations set that might provoke refolding of the amino-terminal domain of the nsP4 RdRp.

Dissociation of the compact form of nsP4 from the nsP1 dodecamer breaks interactions with Arg-85 and Ser-90 (see Compact in **Fig. 9C**). These residues, in turn, initiate a refolding event by forming new interactions with nsP4 (see Extended in **Fig. 9C**). Movement of Arg-85 causes a large change to Gly-83. This residue goes from a *relaxed* conformation, φ, ψ of −65, −46, in the compact state to a *strained* conformation, φ, ψ of −112, −23, in the extended state that can only be accommodated by a glycine residue (see Gly-83 in **Fig. 10C**). Consistent with conservation of this strained conformation is the absolute conservation of this residue (Neff of 1.00 in **Table 3**).

Movement of Ser-90 causes substantial change to nearby residues 91-97. Pro-93, Arg-94, and Ser-97 change conformation in going from compact to extended (**Fig. 10C**). The most striking change occurs at Pro-93. In the compact state, this sidechain has no interactions. In the extended state, Pro-93 is buried in a pocket formed by residues in the 240-250 range (**Fig. 10C**). Pro-93 is not highly conserved and clearly co-varies with residues in the 240-250 range with which it interacts in the extended state (**Supplementary Table S1A**).

Glu-33 also goes from no interaction in the compact state to a well-defined pocket in the extended state (**Fig. 10C**). Glu-33 is quite well conserved (Neff of 1.18 in **Table 3**) but the sequence analysis reveals co-variation with residues 453, 456, and 457 (**Supplementary Table S1A**), residues that the model of the extended conformation has interacting with Glu-33 (**Fig. 10C**).

Together, these results are consistent with both the extended and compact states being evolutionarily conserved across the alphavirus genus.

### Observation of the compact conformation in solution

In parallel to the studies of processed nsP4, we evaluated our ability to purify forms of P34 precursors reported in the past by Lemm and Rice (32). One such precursor, CT50-P34, contains the 50 carboxy-terminal residues of nsP3 linked to nsP4. We purified this precursor (**Fig. 11A**), and it was well behaved enough in solution to perform SAXS (**Supplementary Figs. S16**-**S20**). Surprisingly, the envelope observed for CT50-P34 was consistent with a tetramer of CT50-P34 molecules (**Fig. 11B**). We used the GLOBSYMM module in the ATSAS software suite to fit the observed SAXS electron density with a tetramer formed by monomers in the compact or extended state (69). Clearly, the compact state fit best (compare panel i to panel ii in **Fig. 11C**). A comparison of the biophysical parameters obtained from the SAXS experiments performed on nsP4 and CT50-P34 is presented in **Fig. 11D**. These data provide strong support for the ability of nsP4 to adopt two distinct states, with the compact state exhibiting formation of quaternary structure not observed for the extended state. Together, the SAXS experiments of the two conformational states of the alphavirus RdRp formally adds this protein to the growing list of fold-switching proteins (70,71). In solution, precursor forms of nsP4 may favor the compact conformation while fully the processed form of nsP4 favors the extended conformation. Additional studies will be required to fully understand the structure and biology of the CT50-P34 derivative.

**Figure 11.**
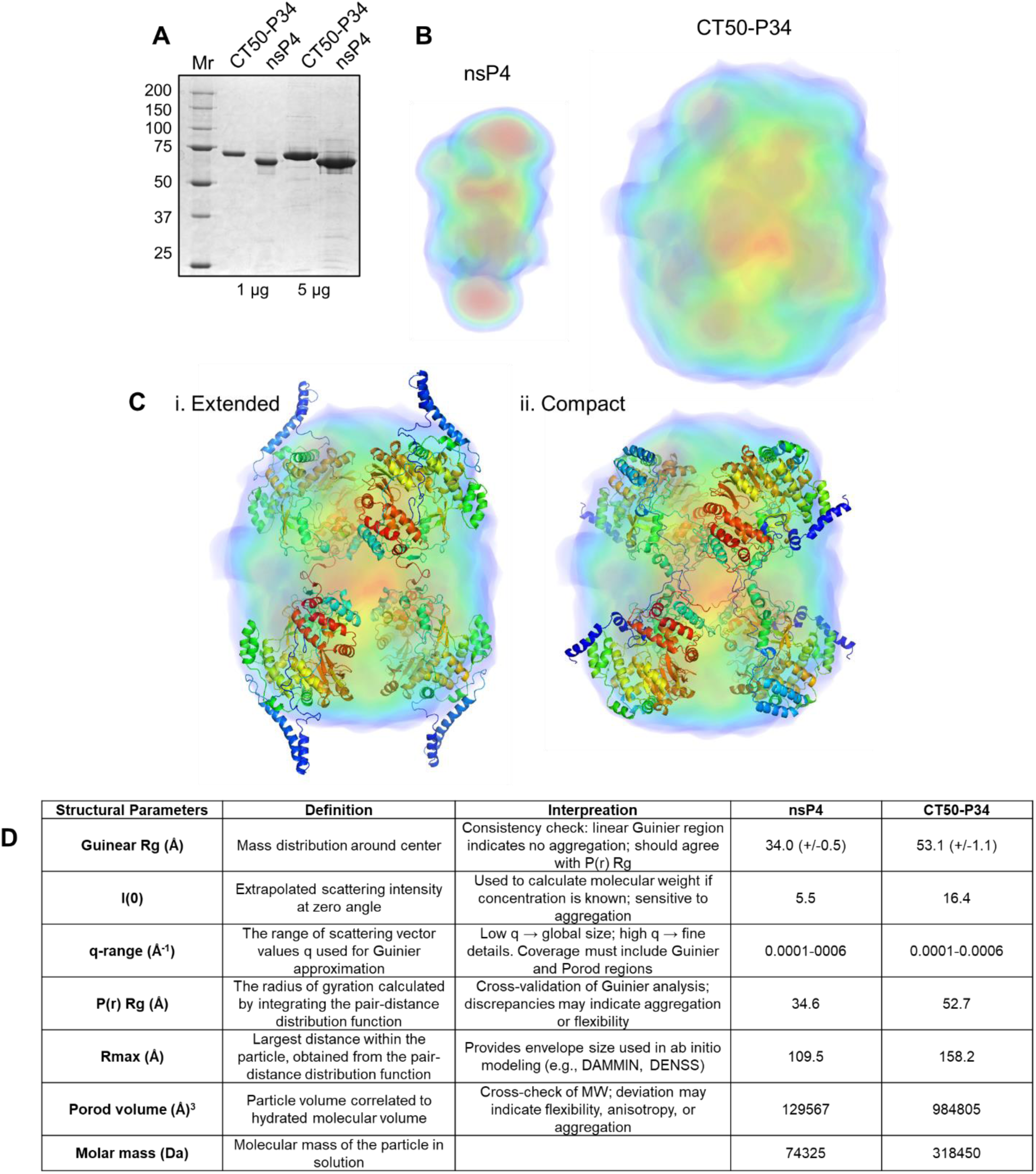
CT50-P34 adopts the compact conformation in solution. (**A**) SDS-PAGE analysis of bacterially expressed and purified CT-50-P34 (75 kDa) and nsP4 (69 kDa). Shown is a 10% polyacrylamide gel with a total of 1 and 5 µg of CT50-P34 and nsP4. Broad-range molecular weight markers (Mr) and corresponding molecular weights are indicated. (**B**) SAXS. SAXS envelope for nsP4 and CT-50-P34. Envelopes are shown to scale. (**C**) GLOBSYMM module in the ATSAS software suite to fit the observed SAXS electron density with a tetramer formed by monomers in the extended or compact state. (**D**) SAXS structural parameters for nsP4 and CT50-P34.

## DISCUSSION

Alphaviruses represent a significant threat to public health. Over the past few years, there have been outbreaks of one alphavirus or another around the globe (72,73), including the United States of America (74). Currently, there are no therapeutics approved to address alphavirus infection. One reason for this circumstance is that only recently has there been a concerted effort to understand alphavirus-encoded enzymes (23,24,75–79). Unlike most other positive-strand RNA viruses impacting human health, a purified alphavirus RNA-dependent RNA polymerase (RdRp) exhibiting robust activity has yet to be reported (21,22,24,25). The RdRp is such a well-established antiviral target (1–3), and there is a clear indication that alphaviruses are susceptible to antiviral ribonucleosides (80,81). The absence of a biochemical system for the alphavirus RdRp prevents use of in vitro approaches to discover potent antiviral ribonucleotides and precludes rigorous interrogation of the mechanism of action.

Alignment of the Sindbis virus RdRp with those of the picornavirus family revealed a unique additional stretch in the amino-terminal domain of approximately 150 amino acids (22). The first 100 amino acids were thought to be disordered (23), and this conclusion was extended to other alphavirus polymerases (23). Interestingly, AlphaFold predictions suggested an ordered structure of the amino-terminal domain, and this predicted conformation was reported to be consistent with the electron density observed in the structure of the O’nyong-nyong virus (ONNV) RdRp bound to the central ring of the Chikungunya virus (CHIKV) nsP1 dodecamer (24). This same study reported very weak (nsP1)_12_-dependent activity of ONNV nsP4 but also demonstrated the absence of polymerase activity for nsP4 alone (24). This observation was consistent with studies of others who suggested the requirement of other non-structural proteins for nsP4 activity (21,82,83).

Because ONNV nsP4 is so well behaved in solution and there had been no biochemical or biophysical analysis of this protein alone (24), the motivation for this study was to fill this gap. The hope was that a low-resolution analysis might inspire hypotheses for the (nsP1)_12_-dependence of the polymerase activity. Consistent with expectations, ONNV nsP4 was easily purified (**Fig. 1**), exhibited secondary structure content consistent with AlphaFold predictions (**Fig. 3**), was monomeric (**Fig. 4**), but lacked the high-affinity RNA-binding activity and polymerase activity observed for other viral RdRps (**Fig. 2**). The story became more intriguing when we performed analytical ultracentrifugation, which suggested an elongated conformation of nsP4 instead of the globular conformation predicted by AlphaFold (**Fig. 5**). The elongated conformation was further supported by the small-angle X-ray scattering (**Fig. 6**) and hydrogen-deuterium exchange mass spectrometry (**Fig. 7**).

Our empirical, biophysical data demonstrate that ONNV nsP4 exists in an *extended* conformation. Such a conformation potentially explains the absence of RNA-binding and RdRp activity for purified nsP4 (**Fig. 2**). Even more intriguing, however, is the fact that AlphaFold did not predict the extended conformation but an alternative, *compact* conformation (24). We suggest that ONNV RdRp, and by analogy other alphavirus RdRps, can adopt two distinct conformations: an active, compact conformation and an inactive, extended conformation.

Transitions from active to inactive conformations are not unprecedented for RdRps of positive-strand RNA viruses. For example, the hepatitis C virus (HCV) RdRp adopts an inactive conformation in solution (84–86). The carboxy-terminal tail of HCV RdRp folds into the RNA-binding channel and interacts with a loop, referred to as the β loop, that is required for initiation of RNA synthesis de novo by the enzyme (87). HCV expresses a polyprotein that is processed post-translationally by its encoded protease. The thought is that a precursor form of the RdRp may be used to establish the complex responsible for initiating genome replication and proteolytic processing, activating the complex. When the RdRp dissociates from the end of the template, it would assume an inactive conformation. Were the RdRp to remain active in the cytoplasm of the cell, the enzyme would bind to cellular RNAs, perturbing their function. Cellular RNAs might also serve as templates leading to production of dsRNA that would, in turn, activate intrinsic defenses of the cell (88).

In picornaviruses, the primary form of the RdRp (also known as 3D) is part of a polyprotein precursor termed P3 that is a fusion of four proteins: 3A, 3B, 3C, and 3D (14).

However, the primary pathway for proteolytic processing of P3 yields two proteins: 3AB and 3CD (14). Despite the presence of the 3D RdRp domain as a part of 3CD, the 3CD protein lacks any detectable RdRp activity (89). The only form of the 3D RdRp that exhibits activity is one with an authentic glycine amino terminus, produced by proteolytic processing of the 3CD protein by the viral protease encoded in the 3C domain (34) The structural basis for proteolytic activation of RdRp activity is known (90). Briefly, formation of the catalytic site requires hydrogen bonding of the amino terminus (NH_3_^+^) with conserved structural motif A, and this interaction can only occur efficiently when a glycine residue is present at the amino terminus. Evolution of such a complex structural mechanism to achieve an active polymerase is not essential. There are countless examples of cellular polymerases for which analogous mechanisms do not exist. As discussed above for HCV, an active RdRp in the cytoplasm of a mammalian cell will be problematic, making control imperative.

The concept of a precursor form of the alphavirus nsP4 serving as the source of the polymerase in cells is an old one (30,31) that no longer appears in most authoritative reviews on togaviruses (91) or alphaviruses (28,29,92). Sindbis virus produces two polyproteins: P123, when processed forms nsP1, nsP2, and nsP3; and P1234, when processed forms P12 and P34 (30). P34 is a fusion of nsP3 and nsP4 and appears to accumulate in cells (30), in contrast to nsP4, which is degraded by the proteasome (93). Hardy and Strauss proposed that P34 might serve as the source of nsP4 for genome replication (30). Later, Lemm and Rice showed that P34 could function in trans in combination with P123 to support genome replication (31). In fact, attaching the carboxy-terminal 50 amino acids of nsP3 to nsP4, referred to as CT50-P34, was able to support genome replication as well as full-length P34 (32). The existence of P34 is not unique to Sindbis virus, as this precursor has been observed for chikungunya virus (94) and likely exists in other alphaviruses.

These forgotten observations of P34 biology inspired us to ask if we could express and purify CT50-P34. Indeed, we could (**Fig. 11A**). Surprisingly, CT50-P34 formed a tetramer in solution based on the SAXS envelope (**Fig. 11B**). In this instance, however, only a tetramer of the compact conformation from AlphaFold fit the envelope (**Fig. 11C**). These studies of CT50-P34 are still in their infancy, and more comprehensive biochemical and biophysical analyses are underway. However, these initial studies provide compelling evidence for the ability of the nsP4 domain to adopt two distinct states.

Our working hypothesis is shown in **Fig. 12**. Precursor forms of nsP4, for example, P1234 and P34, stabilize the nsP4 domain in the compact active formation. The P34 precursor forms a tetramer (**Fig. 11**), but the tetramer will likely exist in equilibrium with species from monomers (**Fig. 12A**) to tetramers. The parameters obtained from the SAXS data make it clear that the equilibrium strongly favors the tetramer (**Fig. 11D**); however, nsP4 monomer binding to the nsp1 dodecamer could shift the equilibrium. An nsP1 dodecamer can only accommodate one nsP4 monomer (**Fig. 12A**), so as more nsP1 dodecamers form, more nsP4 monomers will be recruited from the P34 tetramer. We propose that binding of the precursor form of nsP4 to the nsP1 dodecamer happens before proteolytic cleavage by the nsP2-encoded protease (shown as the intermediate, I, in **Fig. 12A**), because cleavage outside of the nsP1 dodecamer would yield the inactive conformation (**Fig. 12B**). Dissociation of mature, proteolytically cleaved nsP4 from the nsP1 dodecamer would yield the inactive conformation (**Fig. 12B**). Confidence in the evolutionary conservation of this regulatory mechanism is engendered by the fact that nsP1-interacting residues of nsP4 are specifically sequestered in the extended, inactive conformation (**Fig. 12C** and **Figs. 9** and **10**), thus precluding the extended conformation from competing with the precursor forms of nsP4 for binding to the nsP1 dodecamer.

**Figure 12.**
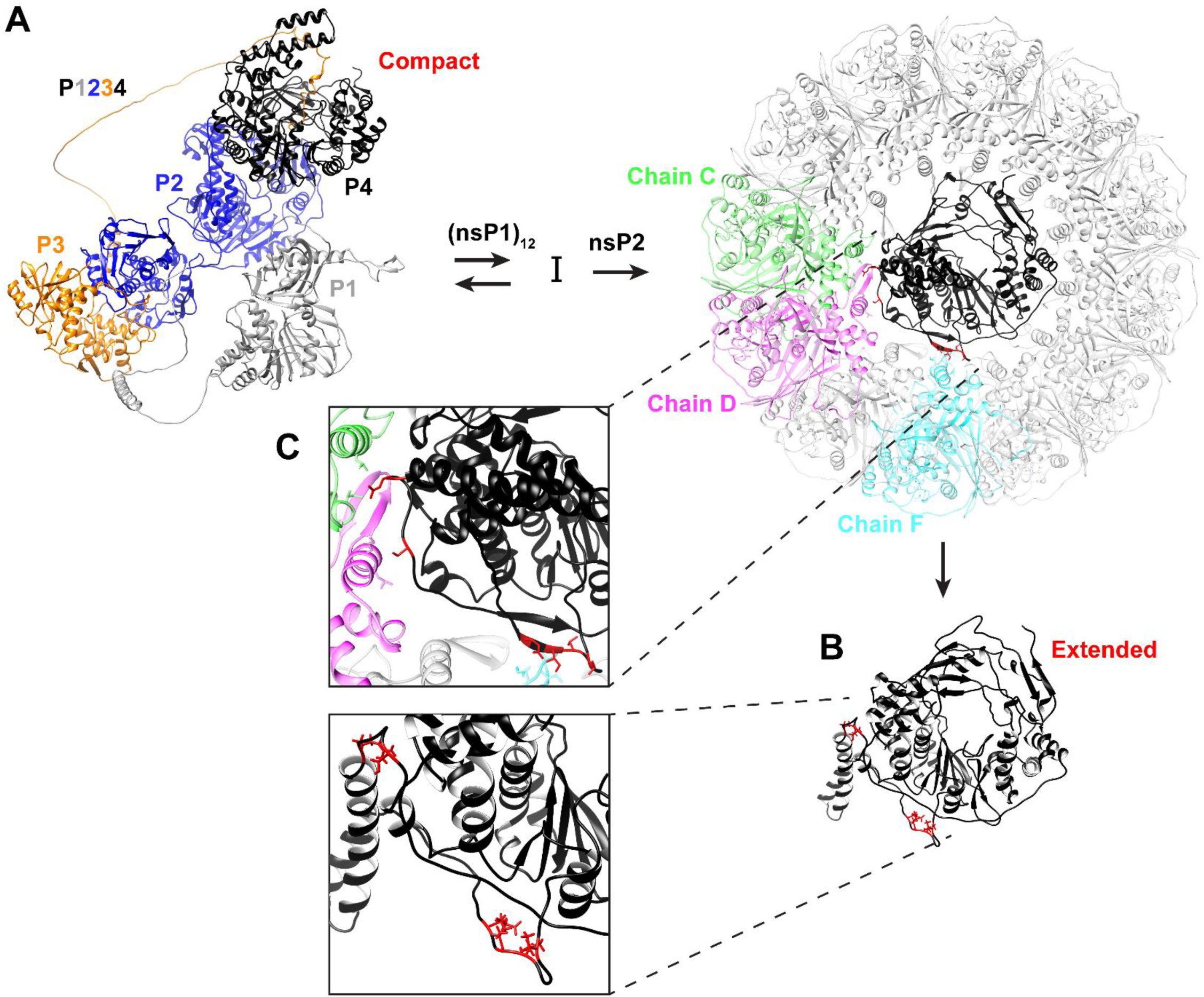
Schematic for P1234 precursor recruitment to the site of replication and fold switching to convert the compact active conformation to an extended inactive conformation. We propose precursor forms adopt the active, compact conformation. At the replication site, proteolytic cleavage would convert the precursor to an active polymerase. Polymerase dissociation upon completion of synthesis would induce fold switching to the inactive, extended state, precluding cytoplasmic activity that would activate intracellular immune responses.

Currently, we are adding the alphavirus nsP4 RdRp to the small but growing list of fold-switching/metamorphic proteins (70,71). Fold-switching proteins are characterized by the unusual ability to adopt two or more distinct, stable folds under near-physiological conditions, with each conformation supporting a different biological function. The switch between folds is typically triggered by environmental cues, including ligand binding or interactions with partner proteins.

In the fold-switching and metamorphic protein literature, proteolytic processing is not considered a mechanism of switching. Here, however, we contend that the fully processed protein interconverts between compact and extended states, with the equilibrium lying overwhelmingly in favor of the inactive extended state. Two observations in the literature are consistent with this interpretation. First, full-length Sindbis virus nsP4 produces minus-strand RNA in vitro in the presence of P123, but only at the limit of assay detection (21).

Second, full-length ONNV nsP4 is able to extend a hairpin RNA template in the presence of the nsP1 dodecamer, but the incomplete conversion of substrate to product over several hours indicates that very little active nsP4 is present (24). For comparison, poliovirus RdRp catalyzes an analogous reaction within milliseconds. Because the extended conformation is not expected to exhibit catalytic activity, the most parsimonious explanation is that a small fraction of nsP4 occupies the compact, active conformation. Future studies will focus on directly detecting and quantifying this equilibrium.

Finally, regardless of whether the transition from the compact to the extended state is a fold switch or a proteolysis-driven conformational change, it should be possible to use time-resolved, structural approaches to monitor the conversion of the precursor form of nsP4 to the mature form by using the nsP2 protease to induce the transition. Such experiments would elucidate the kinetics and structural details of the refolding process and could inspire new strategies to disrupt this remarkable transformation in the alphavirus RdRp.

## ACKNOWLEDGMENTS

The corresponding authors thank members of their teams for their feedback as these studies progressed and for their constructive criticism of the many drafts of the manuscript prior to submission. We also thank Kevin Namitz for assistance in collection and analysis of the analytical ultracentrifugation data. CEC thanks Stan and Thea Sawicki for the many conversations on alphavirus genome replication. His motivation to study nsP4 and many insights conveyed herein would not have happened without those conversations at the annual meetings of the American Society for Virology.

## SUPPLEMENTARY DATA

Supplementary data can be made available upon request.

## CODE AND DATA AVAILABILITY

Data analysis, processing, and visualization was done using a python. These scripts and their associated input and output data are available at: https://github.com/ziul-bio/alpnsp.

