## Supplemental Figures for "A fold switch regulates conformation of an alphavirus virus RNA-dependent RNA polymerase"

### Supplementary Figures, Legends and Tables

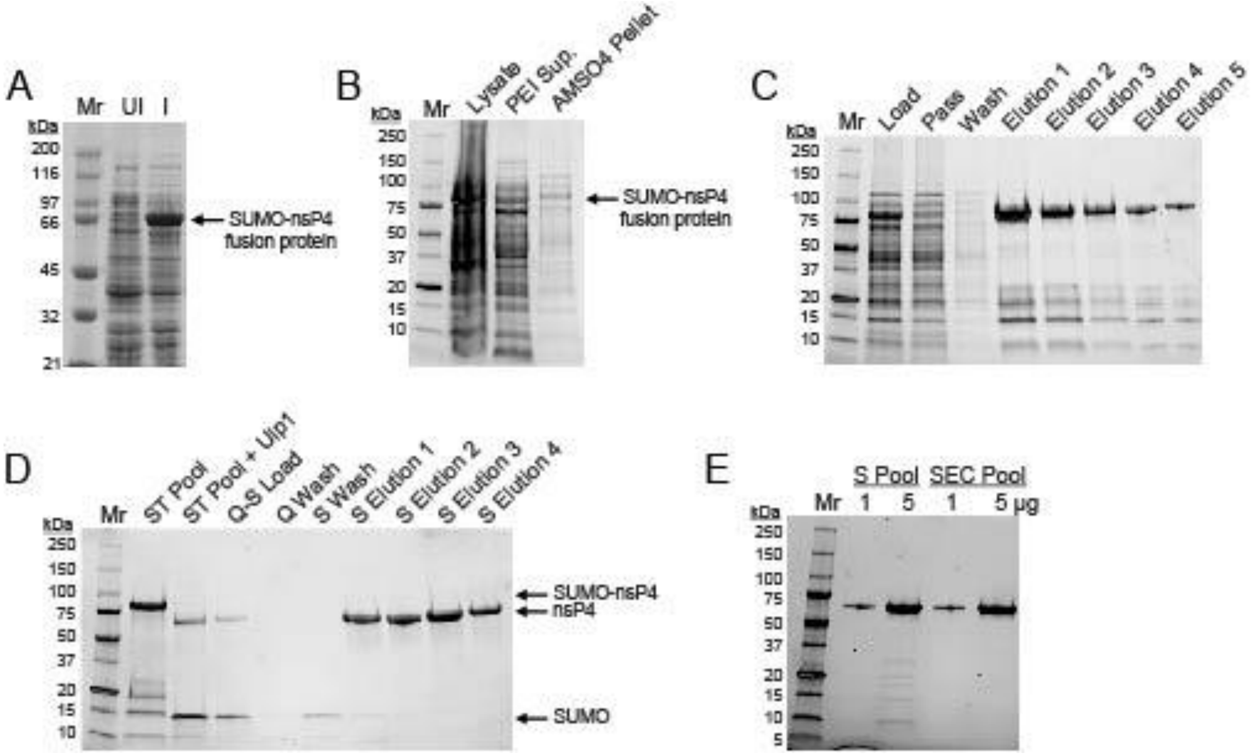


**Supplementary Fig. S1. Expression and purification of ONNV nsP4.** (**A**) SDS-PAGE analysis of bacterially expressed ONNV nsP4 SUMO fusion protein. Shown is a 10% polyacrylamide gel with uninduced and induced samples for the SUMO-ONNV-nsP4 fusion protein. Protein was expressed by Auto-Induction at 15 °C. The position of the induced SUMO-nsP4 fusion protein (81 kDa) is indicated. Broad-range molecular weight markers (Mr) and corresponding molecular weights are indicated. Pre-stained TGX gel was visualized by using a Bio-Rad ChemiDoc MP Imager. (**B-E**) SDS-PAGE analysis of samples from the purification of ONNV nsP4 shown on 4-20% gradient polyacrylamide gels. Panel B: lysate, PEI supernatant, and AMSO4 pellet. Panel C: Ni column column load, Ni column pass, Ni column wash, and Ni column elutions; Panel D: Ni column pooled elutions, Ni column pooled elutions + Ulp1, anion (Q) and cation (S) exchange column purification samples. Cleavage with Ulp1 results in the production of untagged full-length nsP4 protein (69 kDa). The position of nsP4 and cleaved SUMO protein are indicated. Panel E. A total of 1 and 5 µg of nsP4 before (S Pool) and after size exclusion chromatography (SEC Pool).

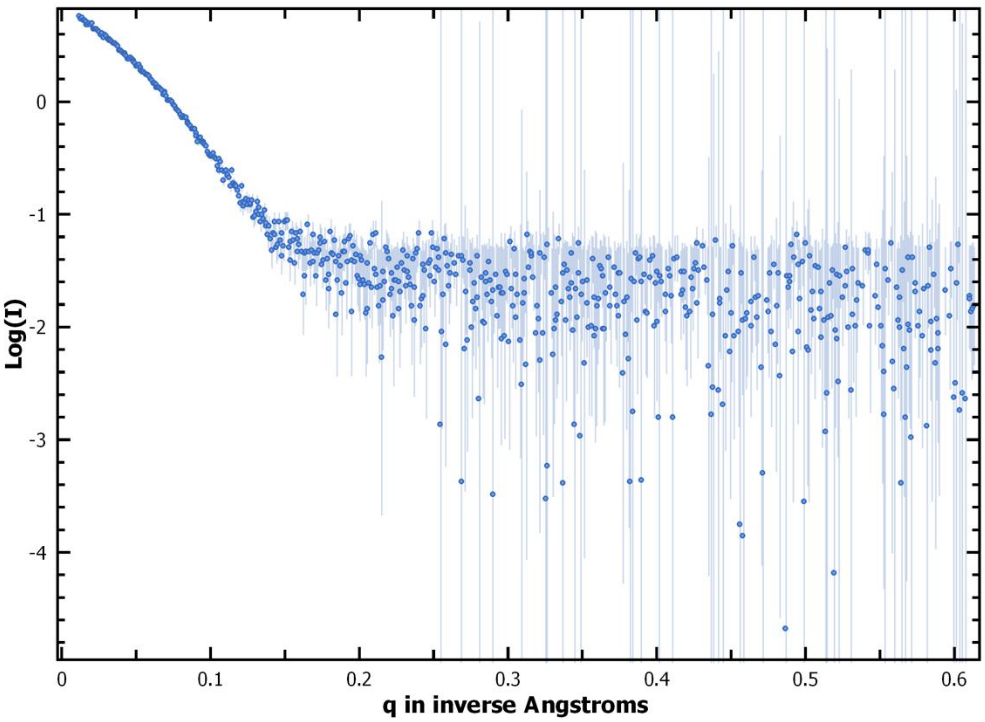


**Supplementary Fig. S2. SAXS raw data.** Shown here is the SAXS raw data for nsP4 protein at 2.9 mg/ml in 25 mM HEPES pH 7.5, 5% Glycerol, 500 mM NaCl, 1 mM TCEP collected on an in-house Rigaku BioSAXS2000$^{nano}$.

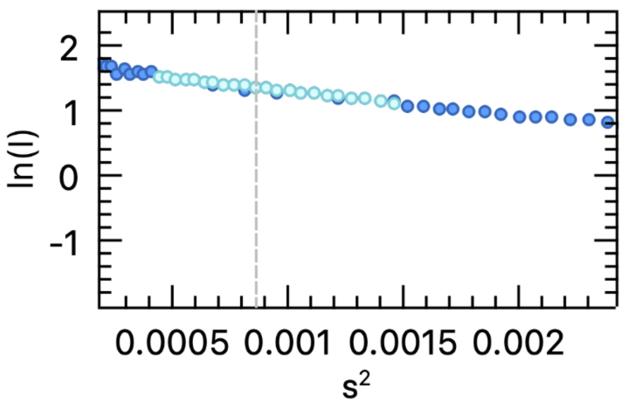


**Supplementary Fig. S3. Guinear plot.** Guinear plot for nsP4 point to a radius of gyration, Rg of 34.0 (+/-0.5)Å which agrees closely with that for the MD model of the extended nsP4 protein with Rg of 34.5Å.

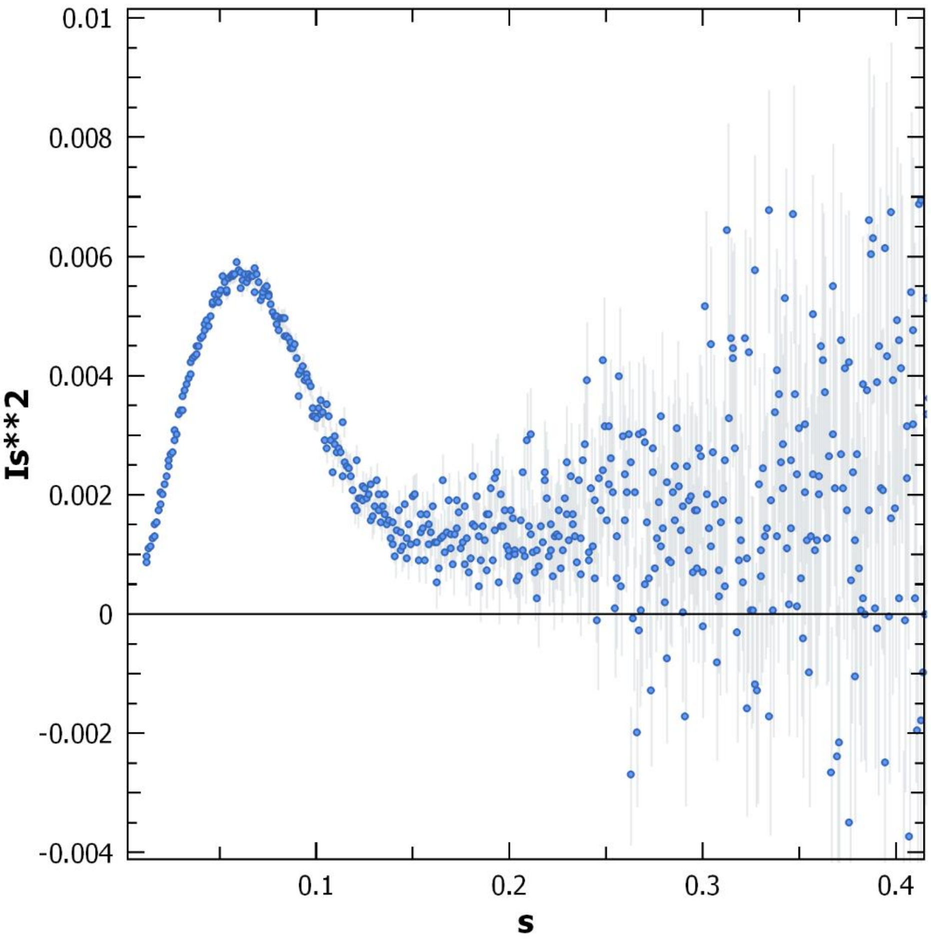


**Supplementary Fig. S4. Kratky plots.** Kratky plot for nsP4 agree with that seen for well folded protein with minimal disorder.

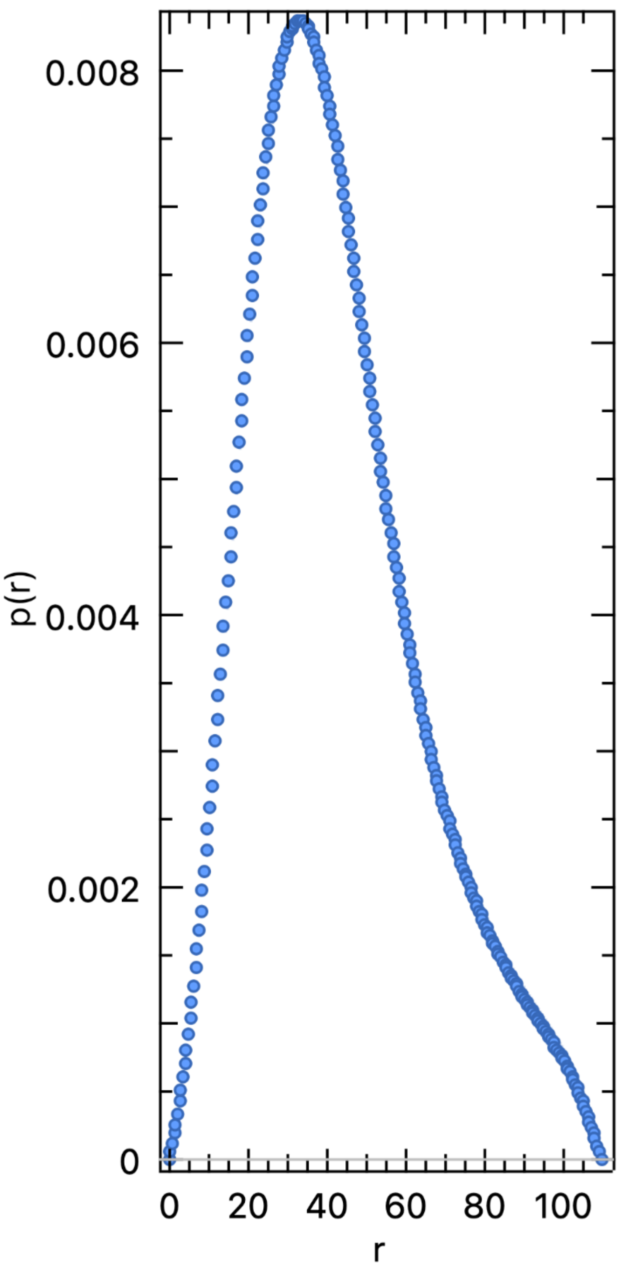


**Supplementary Fig. S5. Pair distance distribution function P(r) function.** Pair distance distribution function P(r) function for the nsP4 peaks at 33.8 Å and in the range of Rg values obtained from the Guinear analysis. The Dmax is around 109 Å in agreement with an extended monomer.

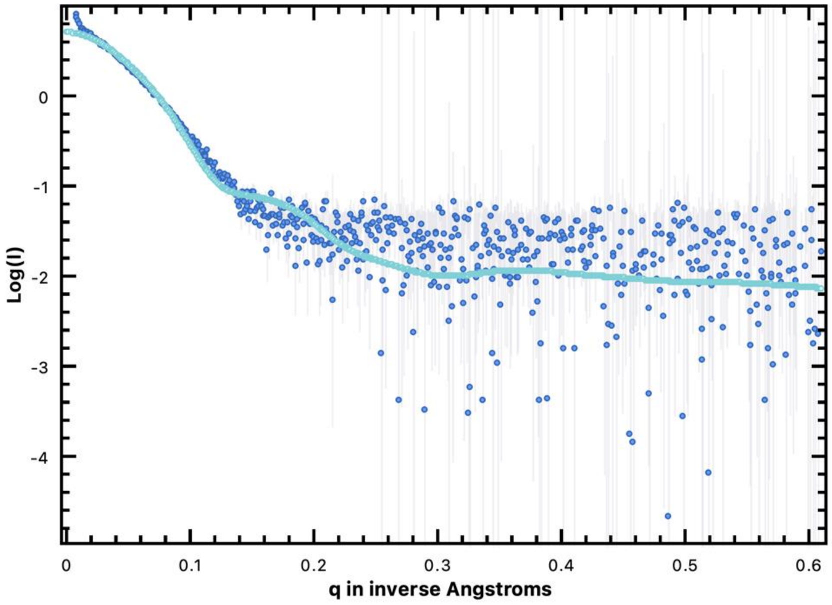


**Supplementary Fig. S6**. **Experimental and calculated SAXS profiles.** Crysol program overlay of the experimental SAXS profile of nsP4 with the calculated SAXS profile as from the Sreflex refined model has a Chi-square fit of 2.6.

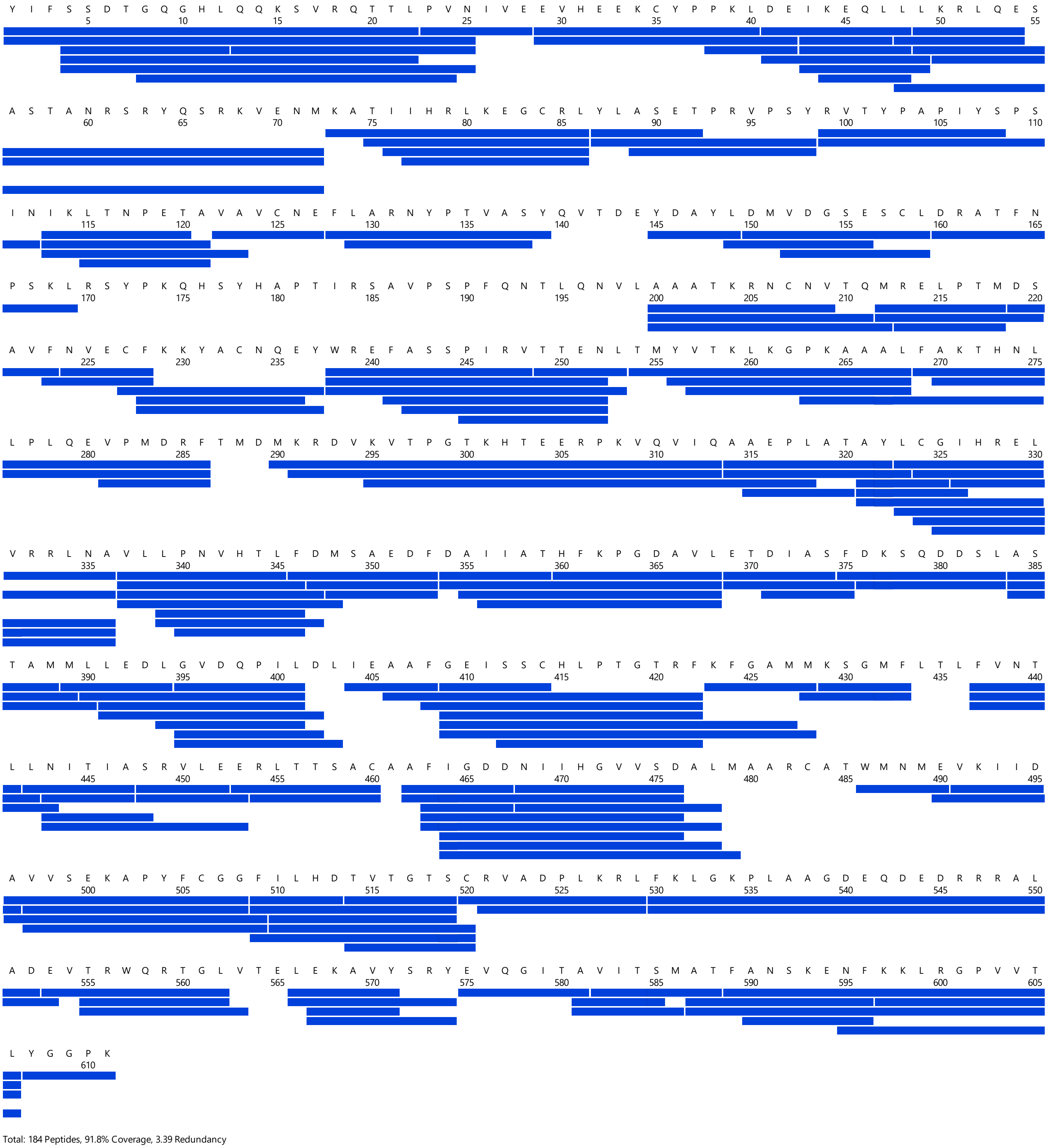

**Supplementary Fig. S7. Sequence coverage map of pepsin proteolyzed peptides of nsP4.** Coverage map showing 184 pepsin proteolyzed peptides for nsP4 spanning 91.8% of the primary sequence with an average redundancy of 3.39.

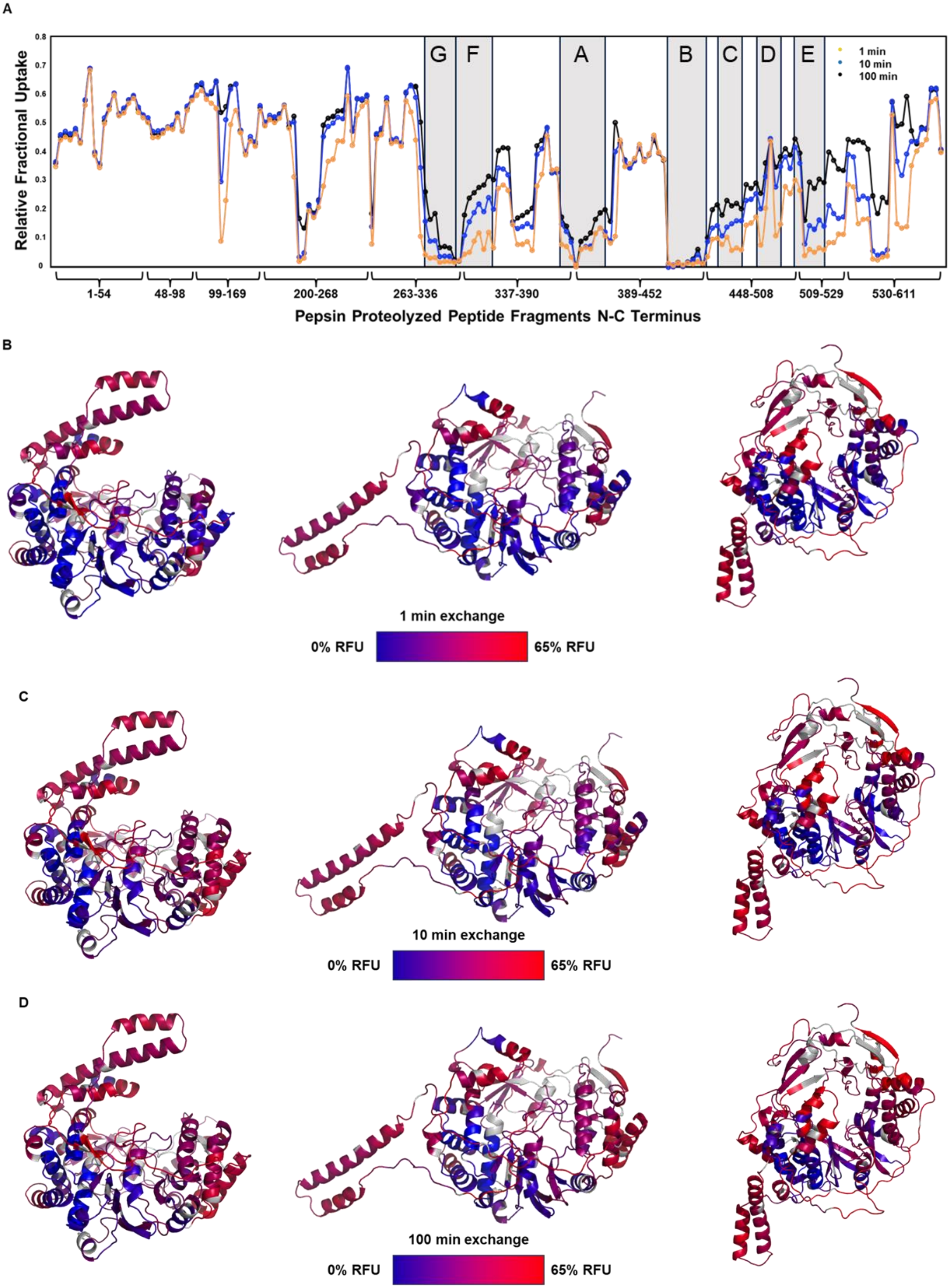
A
Relative Fractional Uptake
G
F
A
B
C
D
E
1 min
10 min
100 min
1-54
48-98
99-169
200-268
263-336
337-390
389-452
448-508
509-529
530-611
Pepsin Proteolyzed Peptide Fragments N-C Terminus
B
1 min exchange
0% RFU
65% RFU
C
10 min exchange
0% RFU
65% RFU
D
100 min exchange
0% RFU
65% RFU

**Supplementary Fig. S8. Relative fractional deuterium uptake of nsP4.** (**A**) Plot of relative fractional uptake after 1 (orange), 10 (blue), and 100 (black) min exchange. Relative fractional uptake corresponds to the ratio of the number of deuterons exchanged to the number of exchangeable amides in a peptide. Each point corresponds to a pepsin proteolyzed peptide fragment plotted from the N to C terminus of the nsP4 sequence. Conserved structural motifs (A-G) are indicated. (**B-D**) Relative fractional deuterium uptake mapped onto the compact (left), SAXS derived extended (middle), and refined extended (right) nsP4 structure after 1 (**A**), 10 (**B**), and 100 (**C**) min exchange. Deuterium uptake is mapped as a gradient with blue representing low relative fractional exchange and red representing high relative fractional exchange. 65% relative fractional uptake corresponds to the maximal deuterium uptake estimated from the highest exchanging peptides after 100 min exchange.

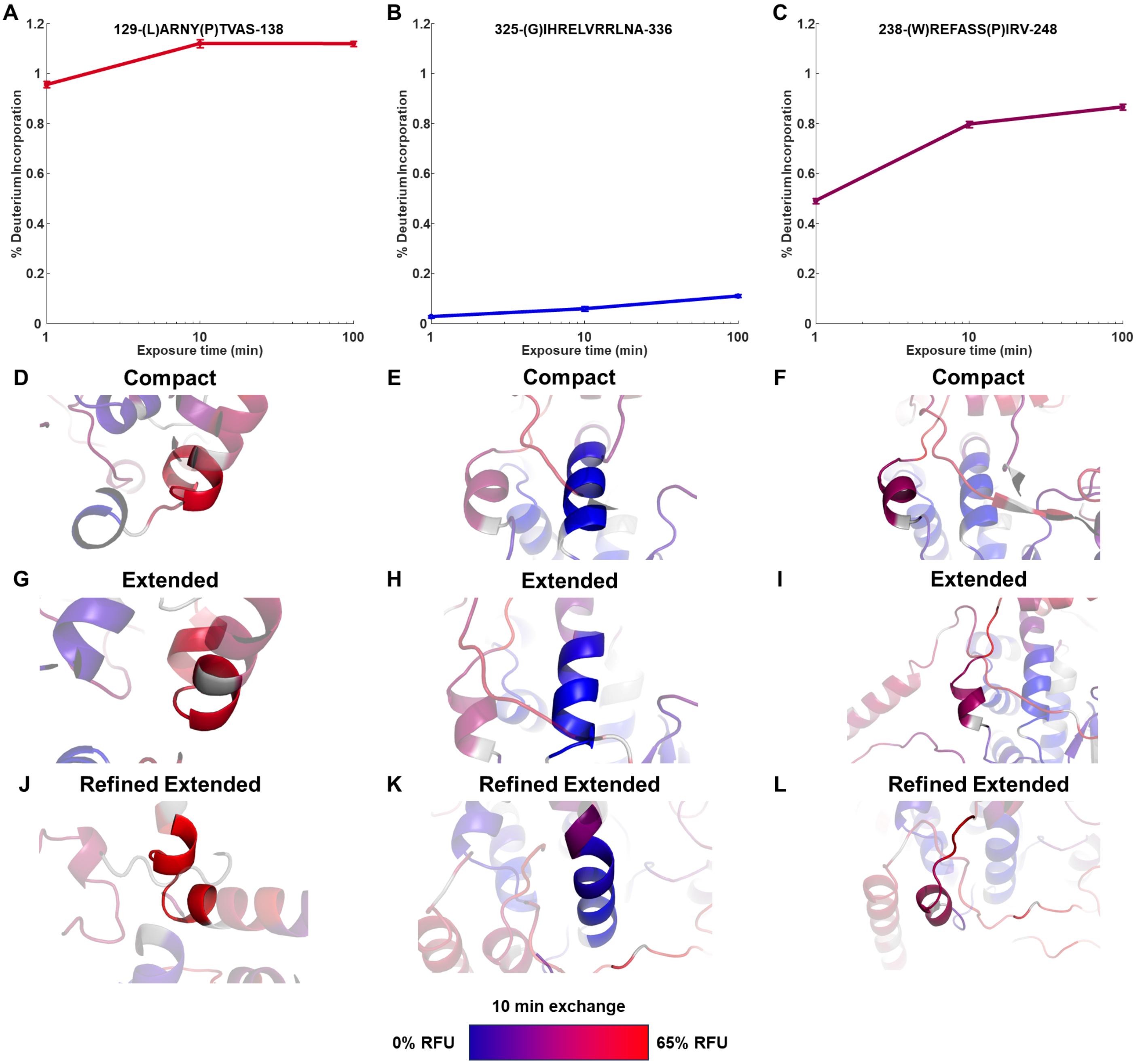


**Supplementary Fig. S9. Deuterium uptake is consistent with common structural features in compact and extended nsP4.** Deuterium uptake plots for representative peptides with common structure in both compact and extended conformations representing (**A**) high, (**B**) low, and (**C**) intermediate exchange. Uptake is plotted as % deuterium incorporation corrected for % deuterium in labelling buffer and average back exchange measured from maximally deuterated samples. Representative peptides highlighted on the compact (**D-F**), extended (**G-I**), and refined extended (**J-L**) structures with relative fractional deuterium uptake after 10 min exchange indicated by a blue-red gradient.

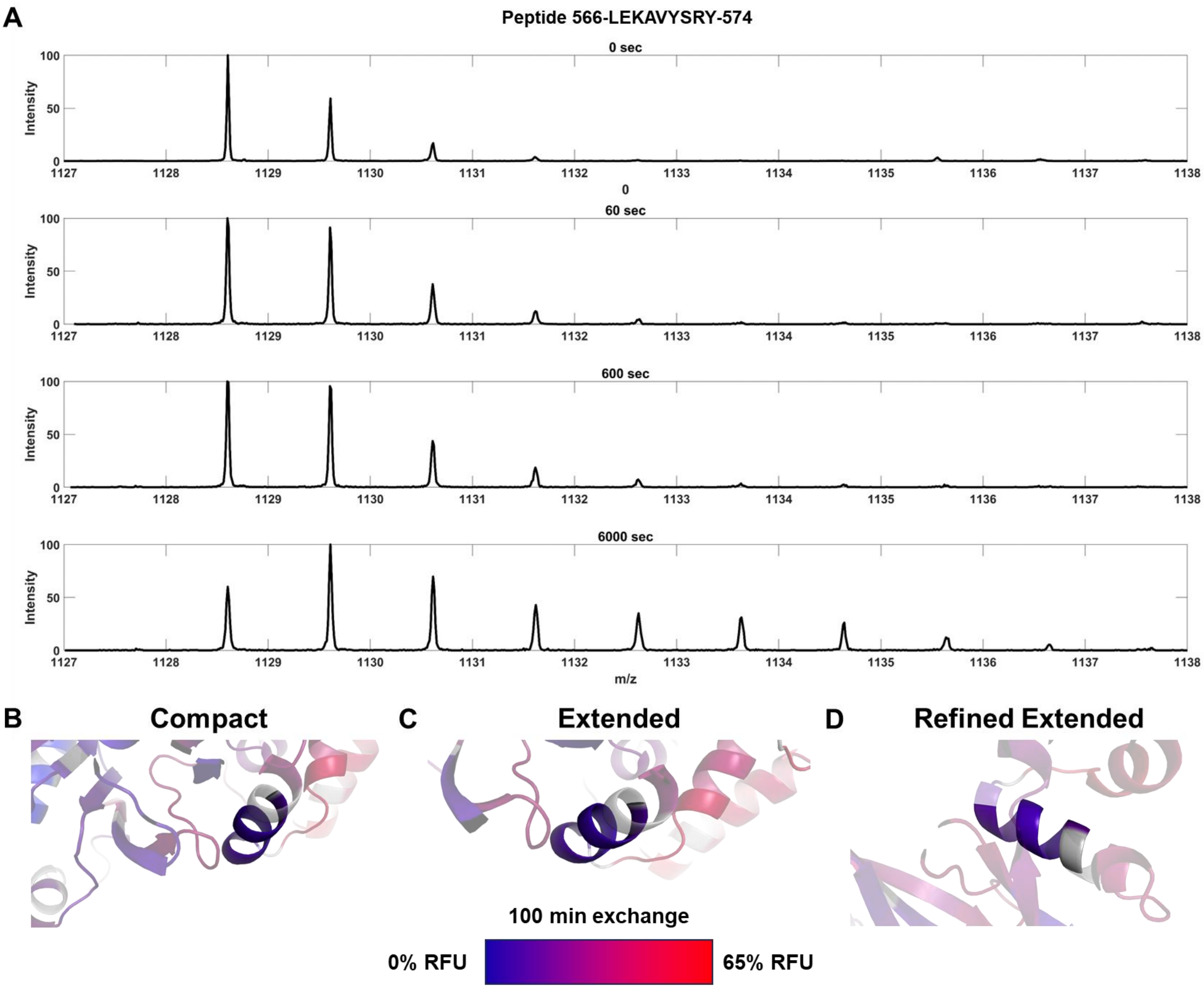


**Supplementary Fig. S10. Only one region of nsP4 shows bimodal hydrogen-deuterium exchange indicating the presence of only one conformation in solution.** (**A**) Mass spectral envelopes for peptide 566-574 for an undeuterated reference state and after 1, 10, and 100 min exchange (top to bottom) revealing bimodal hydrogen-deuterium exchange kinetics. Bimodal hydrogen deuterium exchange kinetics are indicative of two slowly interconverting conformations in solution. No other regions of nsP4 show bimodal hydrogen-deuterium exchange kinetics. Peptide 566-574 is highlighted in the compact (**B**), extended (**C**), and refined extended (**D**) conformation with 10 min relative fractional deuterium uptake indicated by a blue-red gradient.

A

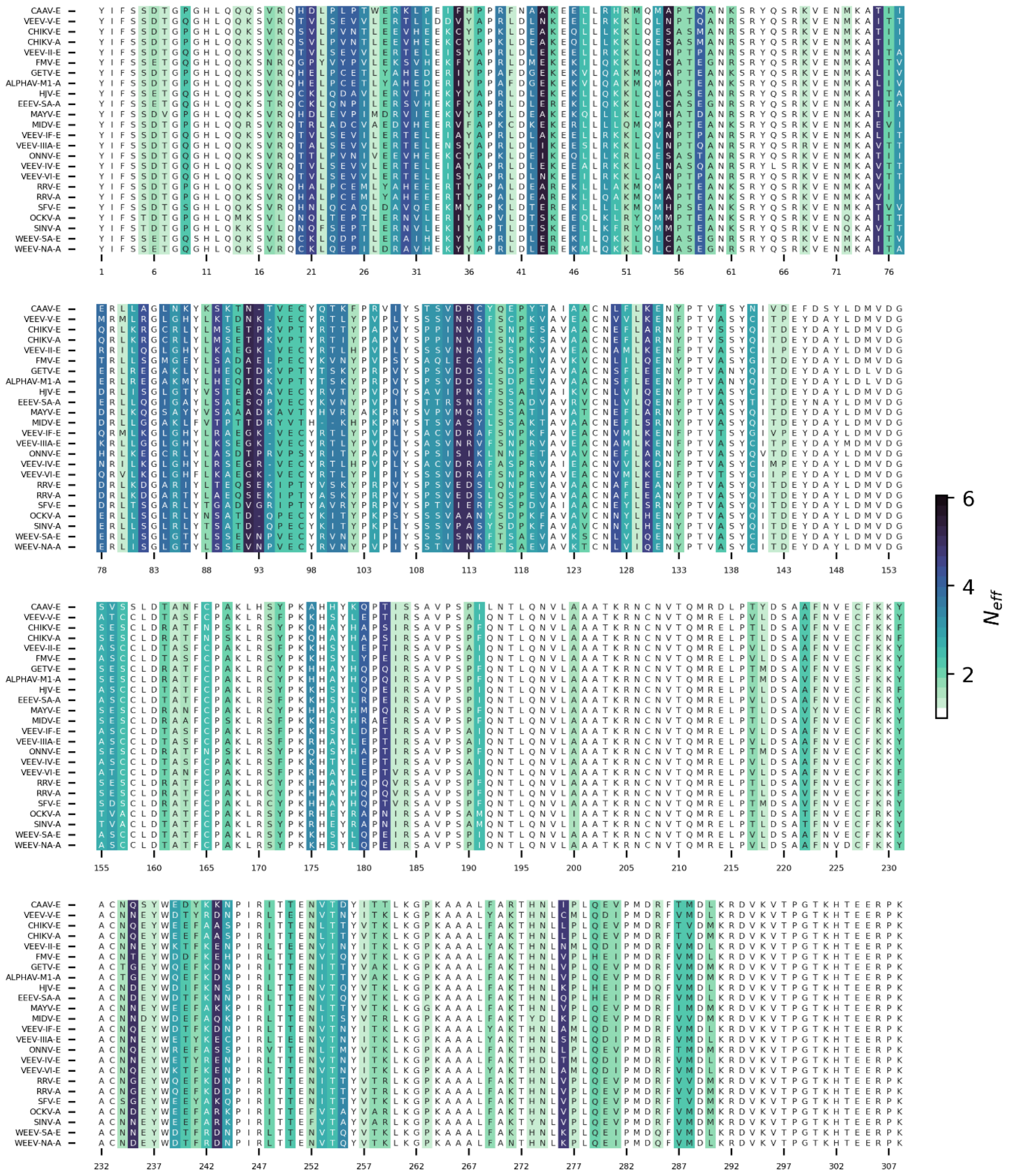
CAAV-E
VEEV-V-E
CHIKV-E
CHIKV-A
VEEV-II-E
FMV-E
GETV-E
ALPHAV-M1-A
HJV-E
EEEV-SA-A
MAYV-E
MIDV-E
VEEV-IF-E
VEEV-IIIA-E
ONNV-E
VEEV-IV-E
VEEV-VI-E
RRV-E
RRV-A
SFV-E
OCKV-A
SINV-A
WEEV-SA-E
WEEV-NA-A
1 6 11 16 21 26 31 36 41 46 51 56 61 66 71 76
78 83 88 93 98 103 108 113 118 123 128 133 138 143 148 153
155 160 165 170 175 180 185 190 195 200 205 210 215 220 225 230
232 237 242 247 252 257 262 267 272 277 282 287 292 297 302 307
6
4
2
$N_{eff}$

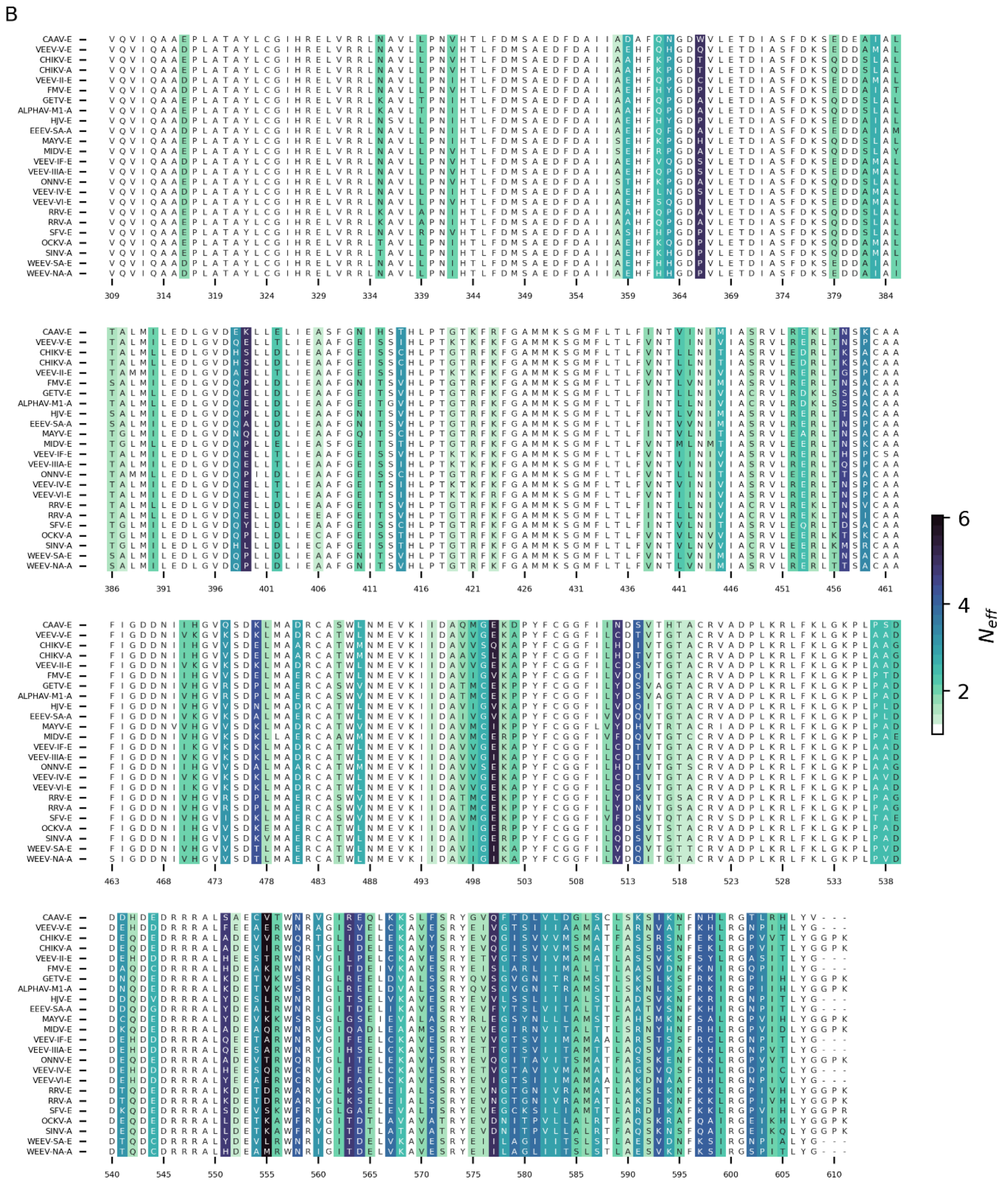


**Supplementary Fig. S11. Conservation of amino acid residues across old- and new-world alphavirus RdRps.** This figure shows a multiple sequence alignment (MSA) of nsP4 proteins from various alphaviruses, divided into two panels for clarity. The alignment visualizes the conservation of amino acid residues across the sequences, with color intensity representing the effective number of sequences contributing to the alignment at each position (*N_eff*). Darker colors indicate lower conservation, while lighter colors indicate higher conservation or lower variability. Panel (A) displays residues 1–307. Alphavirus species are listed on the left, with conserved regions highlighted in the alignment. The heatmap on the right indicates the *N_eff* scale, from low (light) to high (dark) conservation. Panel (B) continues the alignment from Panel (A), covering the C-terminal region of the nsP4 protein, with residues 308–611 displayed, and the same coloring scheme used to represent conservation levels.

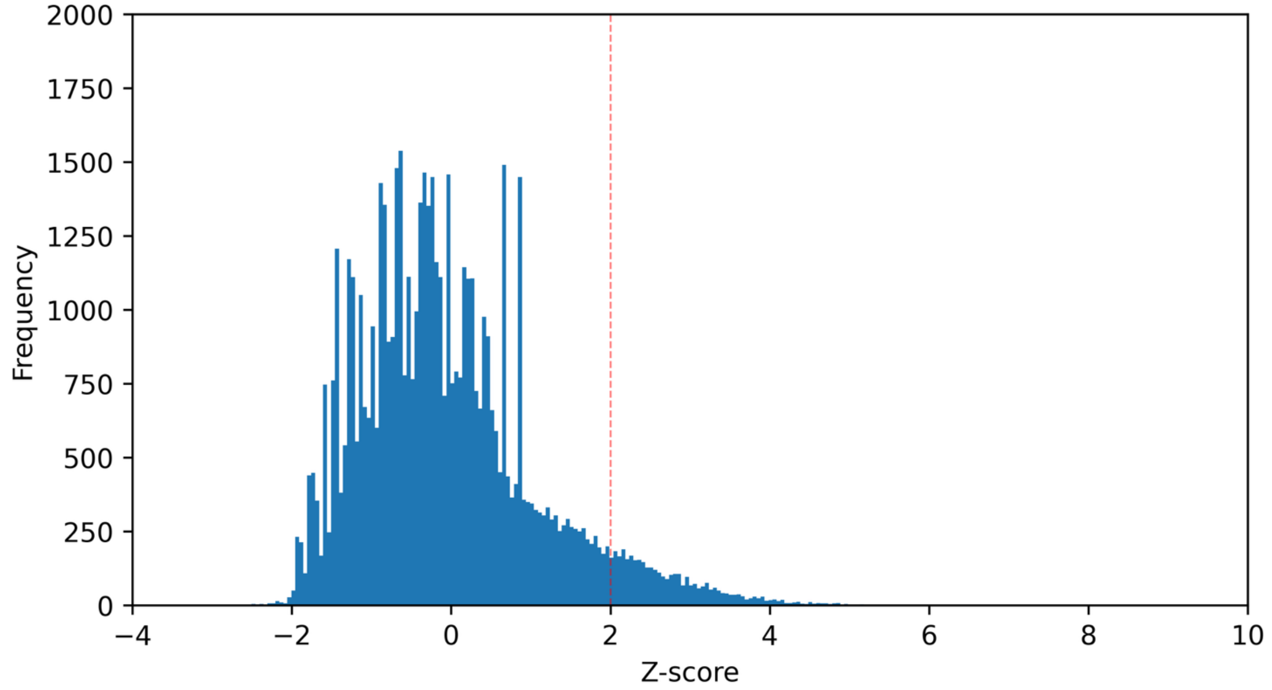


**Supplementary Fig. S12. Distribution of Z-scores for covariance values in the N-terminal region of alphaviruses' nsP4 protein.** This figure analyzes the covariance between residues using Direct Coupling Analysis (DCA), with the covariance values represented as Average Product Correction (APC) and subsequently converted to Z-scores. The histogram displays the frequency distribution of these Z-scores, which highlight the strength of covariation between residues in the RdRp. Most Z-scores are centered around zero, with a significant drop-off as the Z-score increases. We set a threshold of Z-score > 2 to identify significant covariation, as indicated by the distribution tail extending beyond this value, showing the rarity of such Z-scores. This cut-off helps isolate interactions that may be biologically relevant.

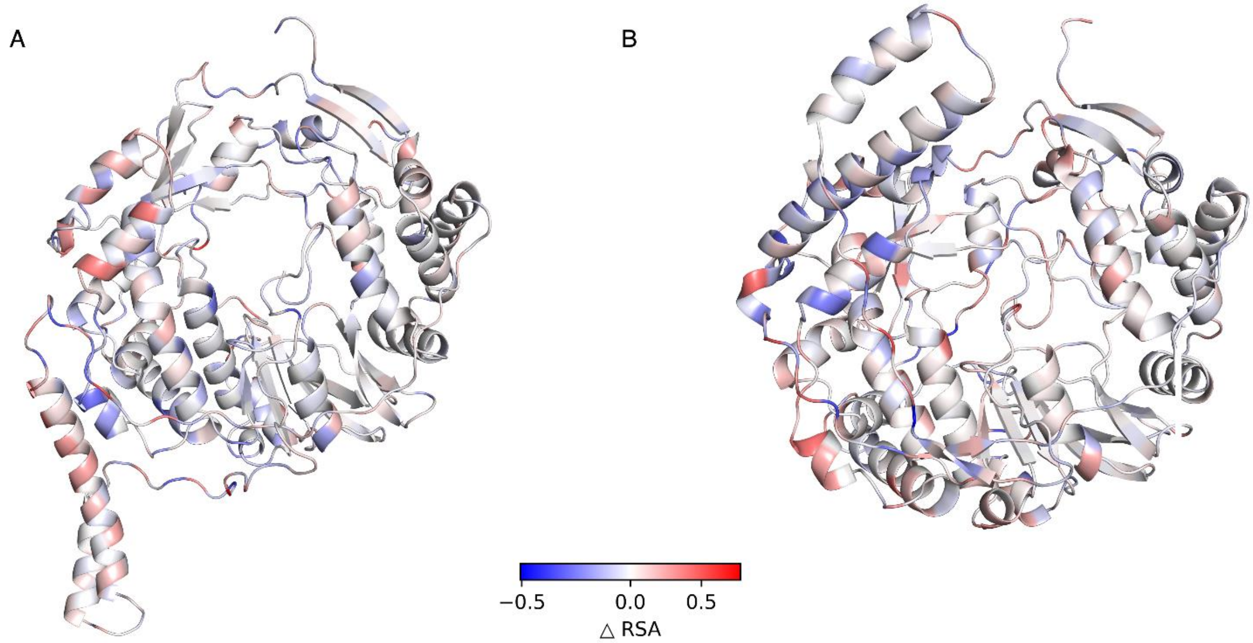


**Supplementary Fig. S13. Structural changes in nsP4 based on differences in relative solvent accessibility (RSA) between the two conformations.** (**A**) Extended conformation of nsP4, with residues colored by RSA differences calculated as *RSA extended - RSA compact*. Blue indicates a decrease in RSA (less exposed in the extended form), red indicates an increase (more exposed), and white indicates no change. (**B**) Compact conformation of nsP4, with residues colored by RSA differences calculated as *RSA compact - RSA extended*, reflecting changes relative to the compact conformation.

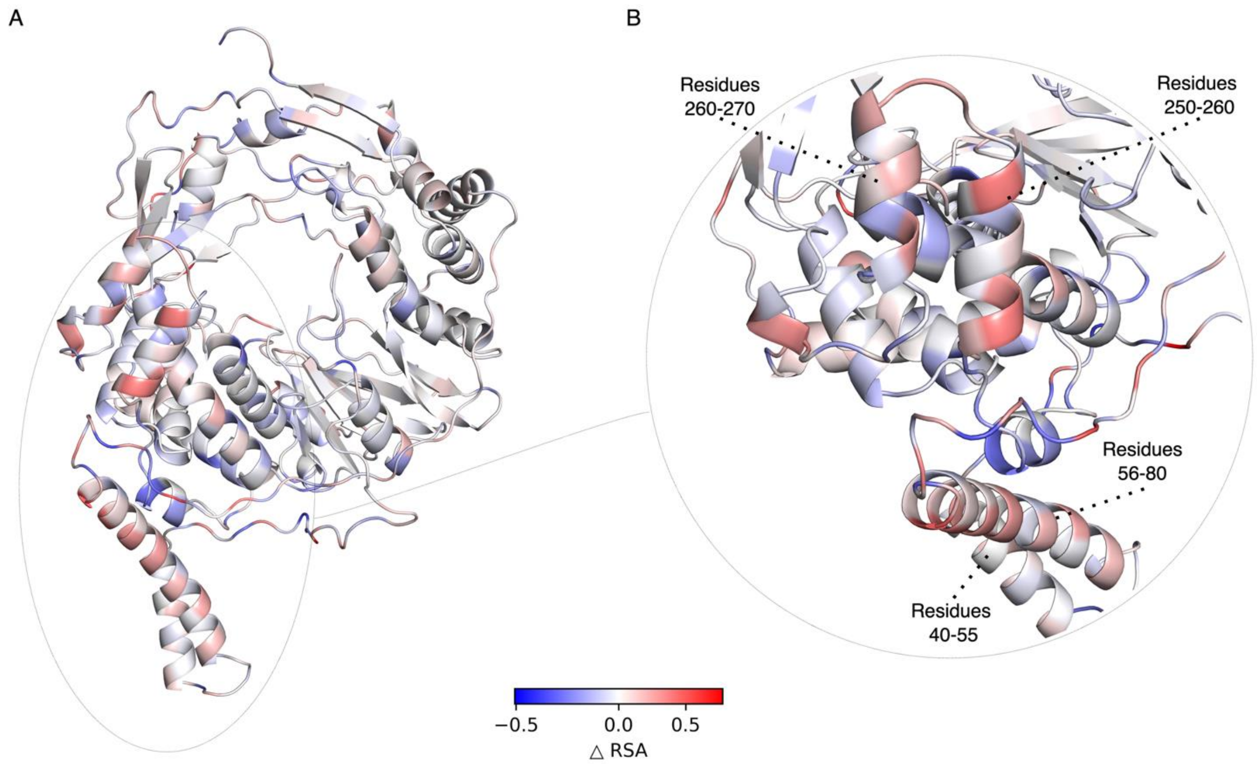


**Supplementary Fig. S14. Structural changes in nsP4 based on differences in relative solvent accessibility (RSA) between the two conformations.** (**A**) Extended conformation of nsP4, with residues colored by RSA differences calculated as *RSA extended - RSA compact*. Blue indicates a decrease in RSA (less exposed in the extended form), red indicates an increase (more exposed), and white indicates no change. (**B**) Compact conformation of nsP4, with residues colored by RSA differences calculated as *RSA compact - RSA extended*, reflecting changes relative to the compact conformation.

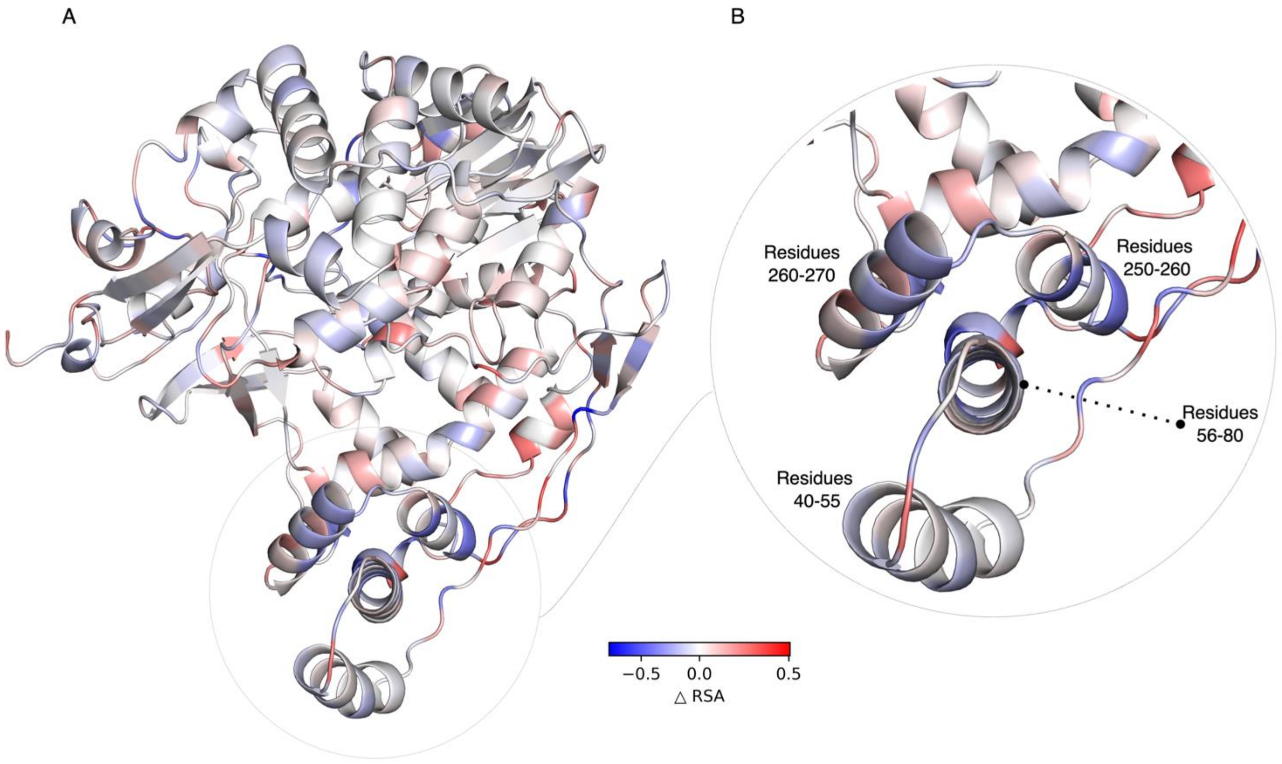


**Supplementary Fig. S15. Visualization of structural changes in the nsP4 protein based on relative difference in RSA between compact and extended conformations.** Panel A displays the entire nsP4 protein, with residues color-coded to represent the relative difference in RSA between the compact and extended forms. Blue indicates a decrease in RSA (indicating less solvent exposure in the compact structure compared to the extended form), red indicates an increase in RSA (indicating more solvent exposure in the compact conformation), and white represents no difference. Panel B offers a zoomed-in view of the N-terminal region, emphasizing residues with the most significant absolute changes in RSA. These highlighted areas correspond to key regions of interest identified in Figure 10, specifically the N-terminal domain (residues 20-90) and the core region (residues 230-270), which exhibit significant structural rearrangements between the two conformations.

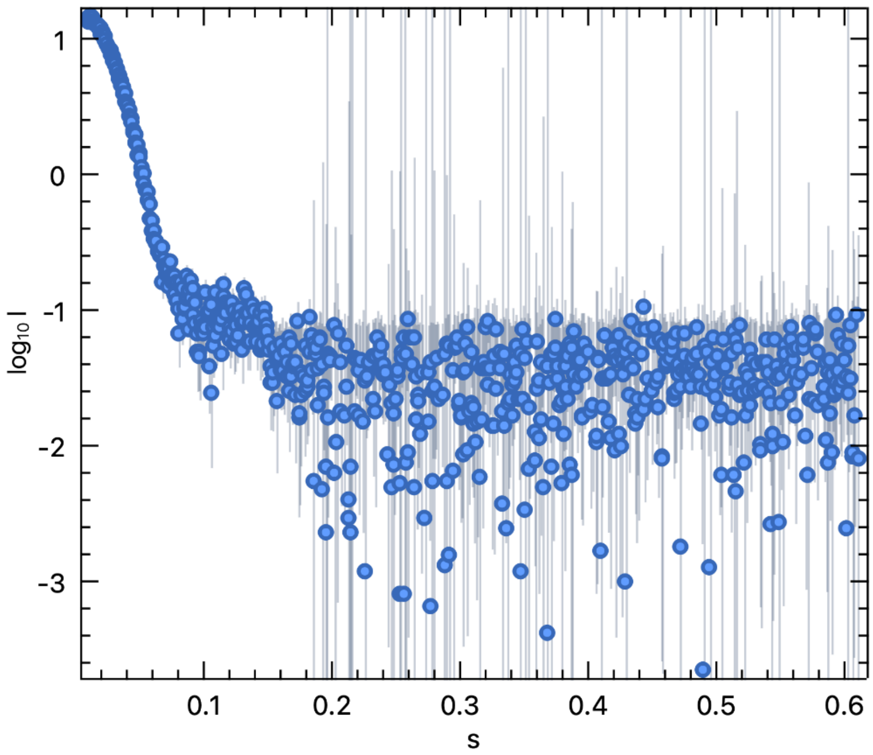


**Supplementary Fig. S16. SAXS raw data for CT50-P34.** Shown here is the SAXS raw data for CT50-P34 at 1.8 mg/ml in 25 mM HEPES pH 7.5, 5% Glycerol, 500 mM NaCl, 1 mM TCEP collected on an in-house Rigaku BioSAXS2000$^{nano}$.

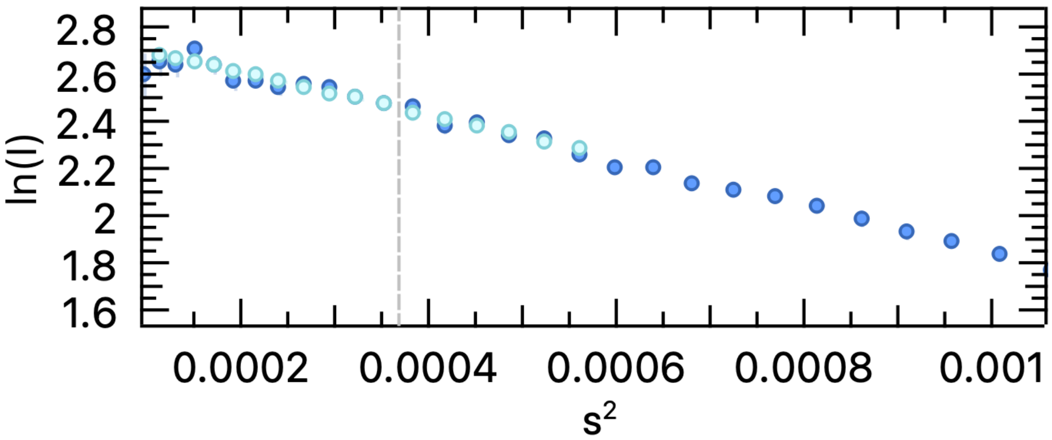


**Supplementary Fig. S17. Guinear plot.** Guinear plot for CT50-P34 point to a radius of gyration, Rg of 53.1 (+/-1.1) Å which agrees closely with a tetramer model of the P34 protein.

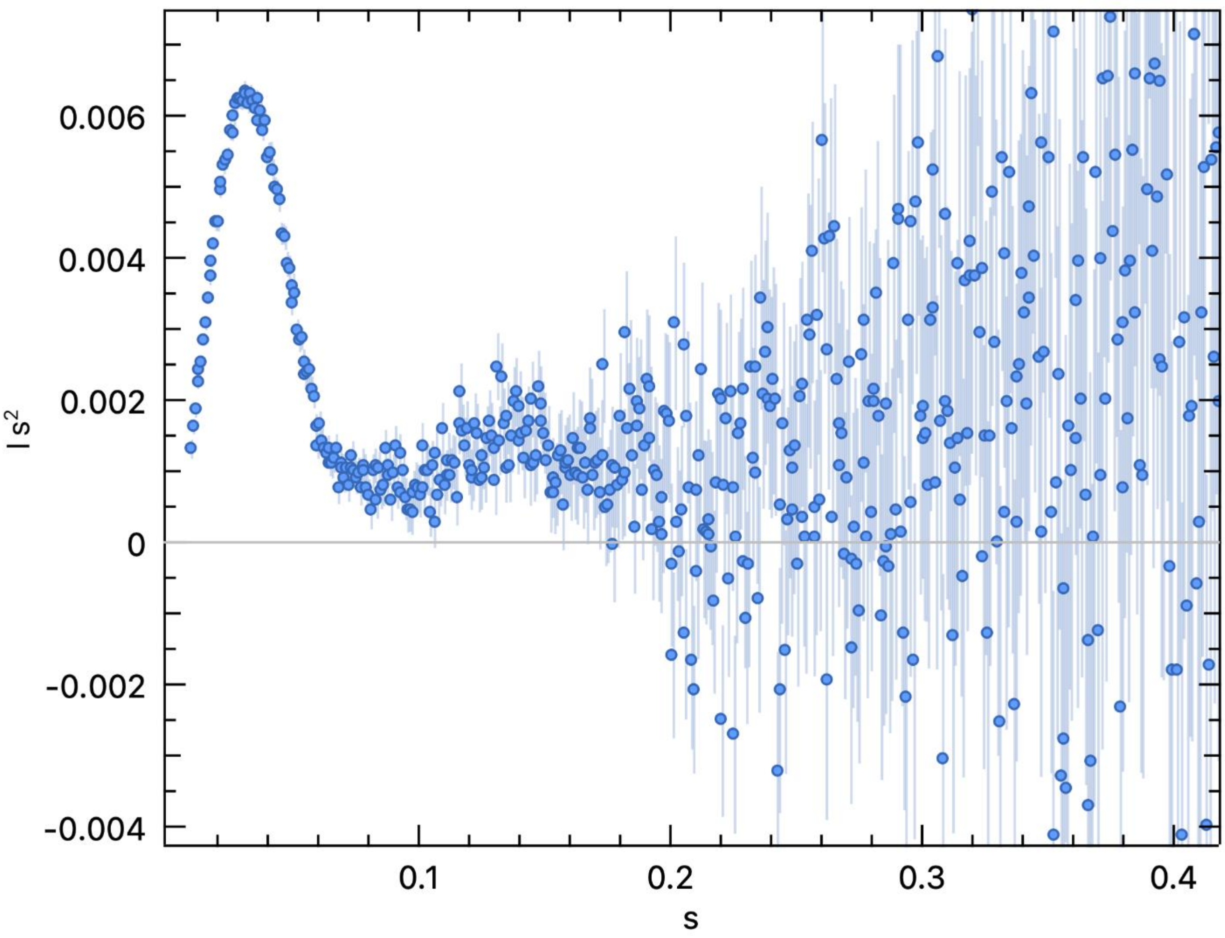


**Supplementary Fig. S18. Kratky plot.** Kratky plot for CT50-P34 agree with that seen for well folded protein with minimal disorder.

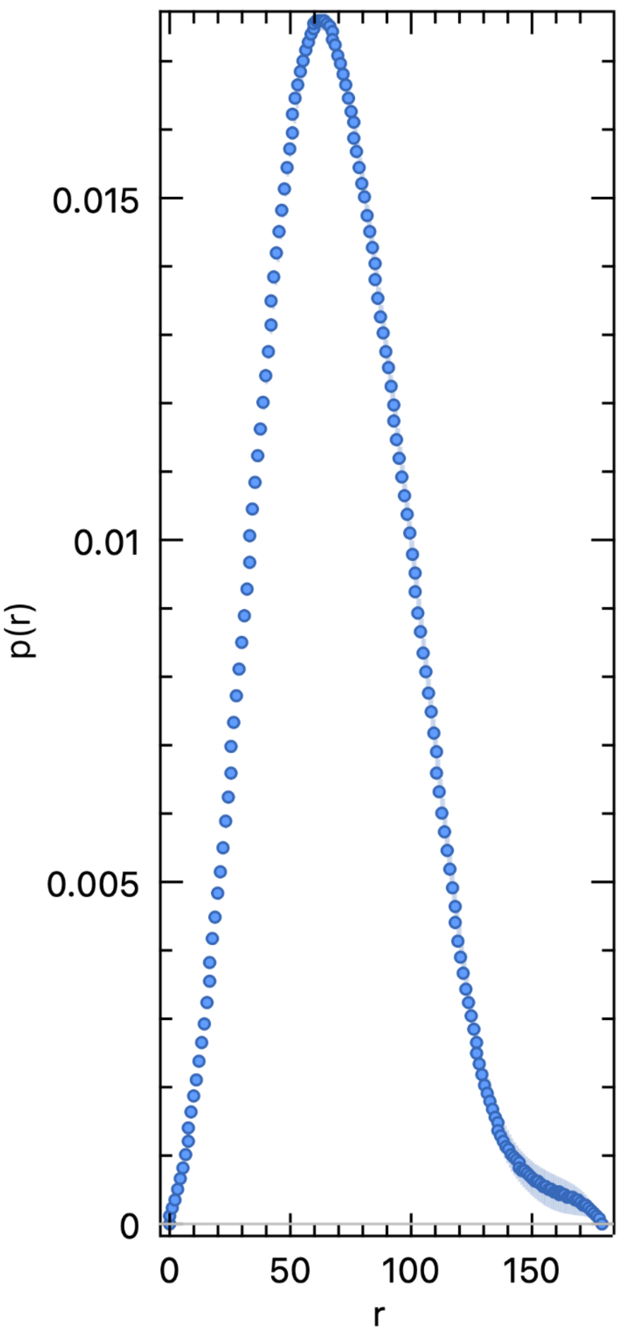


**Supplementary Fig. S19. Pair distance distribution function P(r) function.** Pair distance distribution function P(r) function for the CT50-P34 peaks at 52.7 Å and in the range of Rg values obtained from the Guinear analysis. The Dmax is around 158 Å in agreement with a compact tetramer with Rg and Dmax of 55.9 and 154 Å respectively.

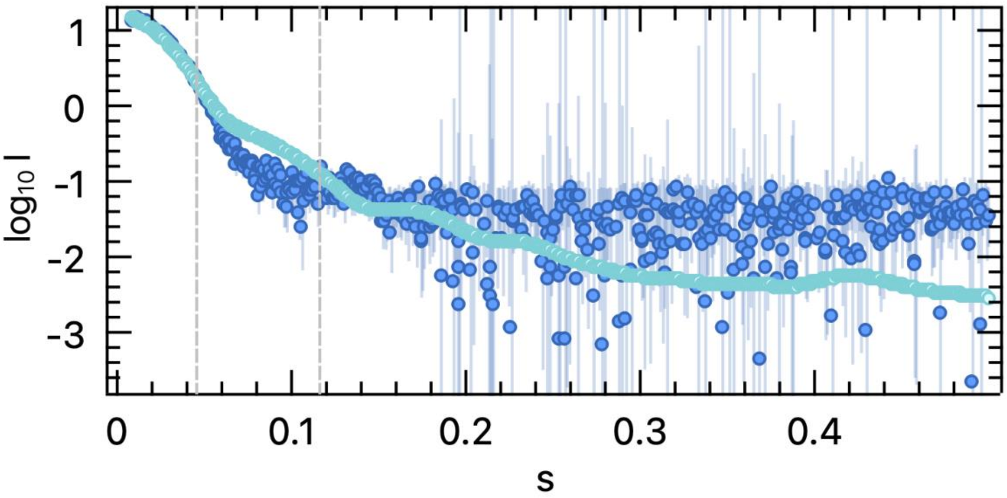


**Supplementary Fig. S20**. **Experimental and calculated SAXS profiles.** Crysol program overlay of the experimental SAXS profile of CT50-P34 with the calculated SAXS profile as from the Globsymm refined compact tetramer model has a Chi-square fit of 6.9.
